# Effects of Host-Dependent Niches and Biotic Constraints on Climate Change Driven Range Shifts in Anemonefish

**DOI:** 10.64898/2026.01.28.702221

**Authors:** Christopher Rauch, Hideyuki Doi

**Author notes:** **Correspondence** Christopher Rauch, Biosphere Informatics, Laboratory, Graduate School of Informatics, Kyoto University, Yoshida-honmachi, Sakyo-ku, Kyoto, 606-8501, Japan. Biosphere Informatics Laboratory, Graduate School of Informatics, Kyoto University, Yoshida-honmachi, Sakyo-ku, Kyoto, 606-8501, Japan (Christopher Rauch). **Funding information**.

## Abstract

We investigated how obligate mutualisms constrain species distributions under climate change, challenging the assumption that biotic interactions are negligible at macro-scales. By integrating host sea anemone distributions into Species Distribution Models for 17 anemonefish species, we found that host availability is a primary determinant of the realised niche, especially for specialists. Under future warming (SSP5-8.5), host immobility creates a biotic constraint, causing fish ranges to lag significantly behind their climatic potential. This mismatch generates over 3.2 million km^2^ of climatically suitable but ecologically inaccessible ocean. Furthermore, specialist anemonefish species with the narrowest niches face the highest climate velocities while being constrained to the most dispersal-limited hosts. These findings indicate that climate-only assessments underestimate extinction risk. Conservation should shift to a host-first management strategy to prevent the collapse of these mutualisms.

**Scientific Significance Statement:** Climate change assessments often assume species can freely track their preferred temperatures, ignoring the critical species they rely on for survival. We demonstrate that for obligate mutualists like anemonefish, the future is defined not just by where they can swim, but by where their host sea anemones can persist. Our models reveal that millions of square kilometers of ocean will become climatically perfect for fish but devoid of the hosts they need to survive. This mechanism of range loss disproportionately threatens specialist species. Our findings highlight the need to prioritise the conservation of immobile partner species, as their failure to migrate effectively traps their mobile symbionts in degrading environments.

## 1 INTRODUCTION

Climate change fundamentally alters the diversity and distribution of species on a global scale Perry et al. (2005); Walther (2010). Across diverse ecosystems, rising temperatures have already triggered large-scale phenological shifts and poleward range contractions, such as the widely documented redistribution of terrestrial butterflies and alpine flora in response to thermal stress Parmesan and Yohe (2003). In marine environments, rising sea surface temperatures and ocean acidification are causing widespread coral bleaching and the degradation of reef habitats Pörtner et al. (2019). These global stressors often combined with local pressures such as coastal development, pollution, and overfishing, exacerbate their habitat degradation and fragmentation Brown et al. (2013); Masucci and Reimer (2019). A large concern for mutualistic systems is the potential for decoupling, where interacting partners respond differently to environmental change, leading to spatial or temporal mismatches that can disrupt the interaction and threaten the persistence of dependent species Walther (2010); Gómez-Ruiz and Lacher Jr. (2019).

*Anemonefish* (Family: *Pomacentridae*) are emblematic coral reef inhabitants known for their mutualistic relationship with host sea anemones. This symbiosis is very important for their survival, as anemonefish depend on anemones for protection, breeding sites, and feeding grounds, while anemones benefit from increased nutrient input and defence against predators Fautin and Allen (1997); Holbrook and Schmitt (2005). However, this partnership is increasingly threatened by global environmental changes, particularly climate change and habitat degradation, which can lead to potential shifts in their species distributions Hobbs et al. (2013); Hayashi and Reimer (2020); Toby Kiers et al. (2010). Specialist species, such as many anemonefish that rely on specific host anemones, would be particularly vulnerable to such decoupling and habitat alterations compared to more generalised species Jiménez et al. (2024a).

The symbiotic relationship between anemonefish and sea anemones is characterised by a spectrum of host species with many anemonefish species relying on a limited range of, or even single, host anemone species Fautin and Allen (1997); Litsios et al. (2012); Gaboriau et al. (2025). This specialisation, coupled with their relatively short pelagic larval dispersal stage Wellington and Victor (1989); Pinsky et al. (2010), makes anemonefish populations particularly sensitive to disruptions in host availability or quality Elliott and Mariscal (2001); Salles et al. (2020). Habitat degradation and fragmentation of anemone distributions due to environmental stressors can therefore directly lead to reduced habitat availability for anemonefish, potentially altering their social structures and decreasing their recruitment success Hobbs et al. (2013); Hayashi et al. (2021). Understanding how these factors, including environmental change, host specificity, and dispersal limitations, collectively influence anemonefish populations and their potential for range shifts is critical for their effective conservation Jiménez et al. (2024b). The Kuroshio Current, a warm northward-flowing ocean current along the Japanese coastline, plays a significant role in facilitating poleward range shifts of marine species, including anemonefish Sudo et al. (2022); Malcolm and Scott (2016); Tanaka et al. (2012). By transporting larvae to higher latitudes, the Kuroshio Current may enable anemonefish and their host anemones to extend their distributions into subtropical and temperate regions Yamano et al. (2011); Perry et al. (2005). Modelling these dynamics is important for predicting how anemonefish populations might shift in response to climate change.

Understanding and predicting the current and future distributions of species is a fundamental goal in ecology and conservation, especially in the context of accelerating global environmental change Pearson and Dawson (2003); Guisan and Zimmermann (2000). Species Distribution Models (SDMs) have emerged as powerful and widely-used tools for this purpose. These correlative models typically relate species occurrence records to a set of environmental predictor variables (e.g. climate and topography) to estimate habitat suitability across a landscape and project it under different scenarios Elith and Leathwick (2009). Furthermore, by stacking individual SDMs, researchers can predict emergent community-level properties, such as patterns of species richness Dubuis et al. (2011); Calabrese et al. (2014); Zurell et al. (2020), providing insights into how entire assemblages might respond to environmental shifts. However, a common limitation of many SDM applications is their primary reliance on abiotic predictors, often overlooking the important role that biotic interactions play in shaping species distributions and community structure Wisz et al. (2013); Godsoe et al. (2017); D’Amen et al. (2018). For species involved in tight ecological relationships, such as obligate mutualisms, the presence and distribution of partner species can be a dominant factor determining their own distribution, potentially overriding purely environmental constraints de Araújo et al. (2014); Guisan et al. (2014). In the context of the anemonefish sea anemone symbiosis, where anemonefish are obligately dependent on their hosts for their survival and reproduction Fautin (1991); Elliott and Mariscal (2001), the distribution of suitable host sea anemones is expected to be a primary biotic predictor of anemonefish occurrence. Incorporating such important biotic factors into SDMs is therefore essential for accurately modelling the distributions of these symbiotic species and for understanding their vulnerability to environmental change Jiménez et al. (2024b); Braun and Lortie (2025).

Given the obligate nature of the anemonefish-sea anemone mutualism and the varying degrees of host specificity observed across species Fautin and Allen (1997); Litsios et al. (2012); Gaboriau et al. (2025), this study aims to quantify the impact of host availability on anemonefish distributions and project how these coupled systems might respond to future climate scenarios. Specifically, we propose three working hypotheses: first, we hypothesise that SDMs incorporating host sea anemone distribution as a biotic predictor alongside environmental variables will exhibit significantly higher predictive performance for anemonefish than models based solely on environmental predictors Jiménez et al. (2024b). We predict this gain in accuracy will be asymmetric, with specialist species showing a greater dependency on biotic predictors than generalists, clarifying the role of the host in defining the realised niche. Second, we hypothesise that host immobility will act as a biotic constraint on climate-driven range expansion. We predict that the realised range shifts of anemonefish (constrained by hosts) will lag behind their potential climatic shifts. We further predict that this constraint will be ubiquitous but significantly stronger in specialist species, who lack alternative partners to facilitate colonisation of new frontiers. finally, we hypothesise that species with narrower realised environmental niches (specialists) will be projected to undergo larger geographical displacements to track suitable climate conditions than broad-niche generalists, effectively subjecting them to higher climate velocity pressure. By testing these hypotheses, this study seeks to advance the understanding of the ecological and biogeographical consequences of obligate mutualism under future climate conditions.

## 2 METHODS

Our methods outline the steps taken to collect species occurrence data, prepare environmental predictor variables, build and evaluate the SDMs, and perform statistical analyses to compare the environmental drivers of both communities. All analyses were performed using R version 4.3.1 R Core Team (2025).

### 2.1 Study Area and Species Occurrence Data

Anemonefish live in social groups and have a symbiotic relationship with ten known species of host sea anemones. These host sea anemones inhabit the tropical Indian Elliott and Mariscal (2001) and western Pacific Oceans Fautin and Allen (1997), with the highest species diversity located in the Indo-Malay archipelago. To capture potential polward range shifts driven by climate change, the study extent was defined to include not only the core tropical Indo-Pacific but also adjacent warm-temperate expansion zones, such as the Southeast Australian Shelf and Northern New Zealand, where the tropicalisation of temperate reefs is increasingly observed Vergés et al. (2014). The final study area is shown in figure 1.

**FIGURE 1.**
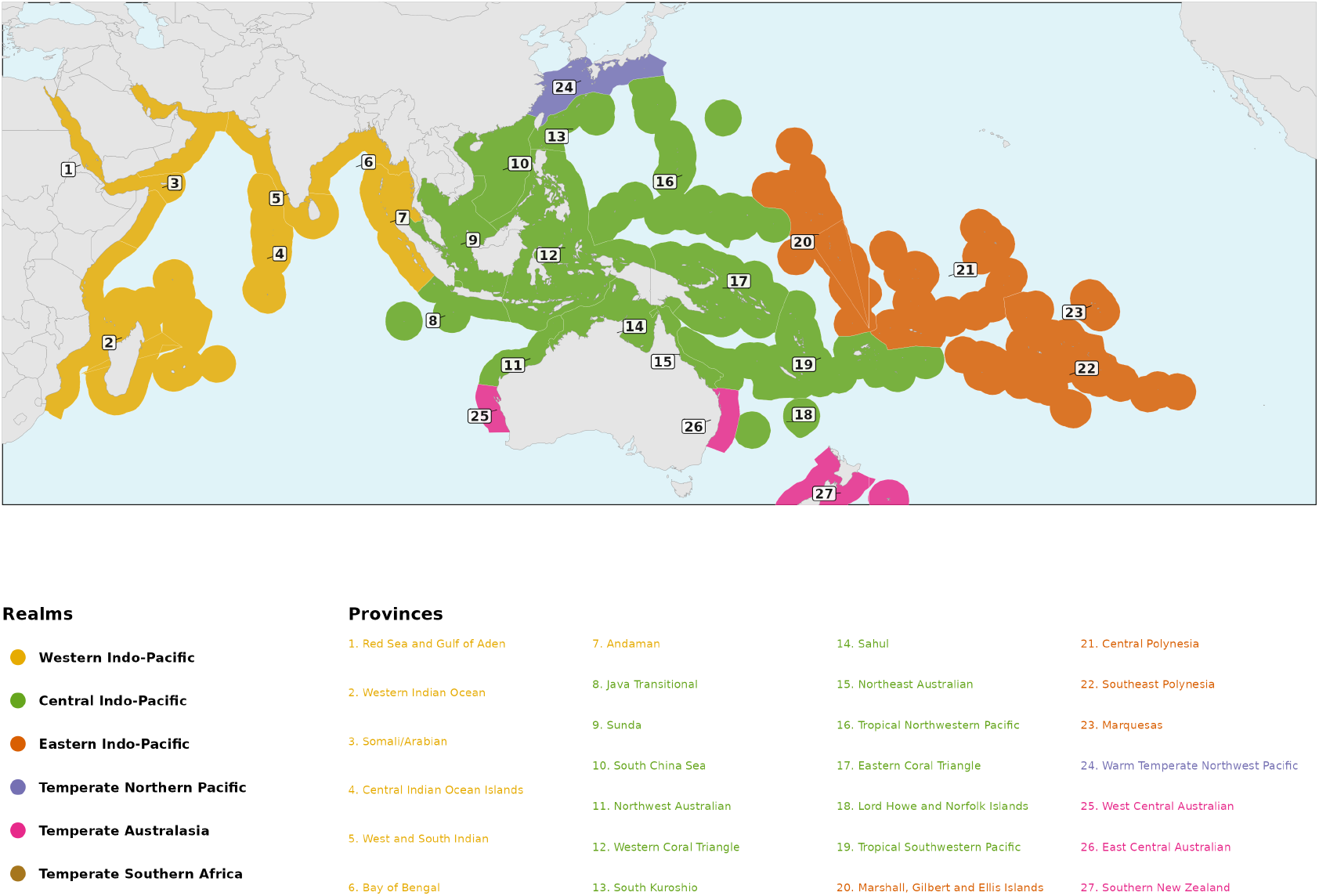
Study extent defined by Marine Ecoregions of the World (MEOW) provinces. Colours indicate biogeographic realms. The study area includes core tropical provinces and temperate expansion zones (e.g., Southeast Australia, Northern New Zealand) to capture potential range shifts.

Occurrence records for sea anemone and anemonefish were collected from public online databases including the Global Biodiversity Information Facility (GBIF) GBIF.Org User (2025) and the Ocean Biogeographic Information System (OBIS) OBIS using the `rgbif` Chamberlain et al. (2022) and `robis` Whiting et al. (2016) packages, respectively. To ensure data quality, raw records underwent a cleaning protocol using the `CoordinateCleaner` package Zizka et al. (2019). We excluded records that were: (1) fossils or aquarium specimens; (2) located on land or at depths > 200 m; (3) collected prior to 1970, to align with the temporal coverage of environmental predictors; or (4) flagged with high coordinate uncertainty (> 10 km). Remaining records were spatially filtered to the defined Marine Ecoregions of the World (MEOW) study extent Spalding et al. (2007) and thinned to a single unique occurrence per ∼9 km grid cell to mitigate sampling bias.

### 2.2 Environmental Predictor Variables

To characterise the abiotic niche, eight bioclimatic variables were sourced from the Bio-ORACLE v2.0 dataset Tyberghein et al. (2012); Assis et al. (2018, 2024), representing key gradients in temperature, salinity, and nutrients (Table 1). Data was downloaded at a spatial resolution of 5 arc-minutes (∼9.2 km at the equator).

**TABLE 1.** Bioclimatic predictor variables sourced from Bio-ORACLE v2.0 used in the Principal Component Analysis. Variables were selected to represent key physiological constraints on anemonefish and host sea anemones.

| Variable Code | Description |
| --- | --- |
| Temperature |  |
| SST Mean | Mean Sea Surface Temperature (°C) |
| SST Range | Range of Sea Surface Temperature (°C) |
| Light |  |
| PAR Mean | Photosynthetically Active Radiation |
| Chemistry |  |
| Salinity Mean | Mean Salinity (Practical Salinity Scale) |
| pH | Mean pH |
| Nutrients |  |
| Nitrate | Nitrate Concentration ($mol \cdot m^{-3}$) |
| Phosphate | Phosphate Concentration ($mol \cdot m^{-3}$) |
| Silicate | Silicate Concentration ($mol \cdot m^{-3}$) |

Given the high multicollinearity inherent in bioclimatic data, Principal Component Analysis (PCA) was employed to reduce dimensionality while retaining environmental information. We retained the first four Principal Components (PC1–PC4) based on a consensus of methods. While the strict Kaiser-Guttman criterion (Eigenvalue > 1) suggests retaining only the first three axes, PC4 exhibited an eigenvalue of 0.87, above the modified Jolliffe criterion of 0.7, which is preferred to prevent the exclusion of ecologically relevant gradients Kaiser (1960); Jolliffe (1972). Retaining PC4 increased the cumulative explained variance from 77.9% to 88.8%, providing a comprehensive representation of the climatic niche (see Supporting Information S1, figure S1). PC5 (Eigenvalue = 0.53) was discarded as noise. The interpretation of these synthetic variables is visualised in figure 2. PC1 represents a primary gradient of surface temperature and light (Photosynthetically Active Radiation, PAR), essentially capturing the latitudinal gradient from the tropics to the temperate expansion zones. PC2 captures gradients in nutrients (Nitrate) and depth-specific temperature ranges, differentiating nutrient-rich upwelling zones from oligotrophic reef flats.

**FIGURE 2.**
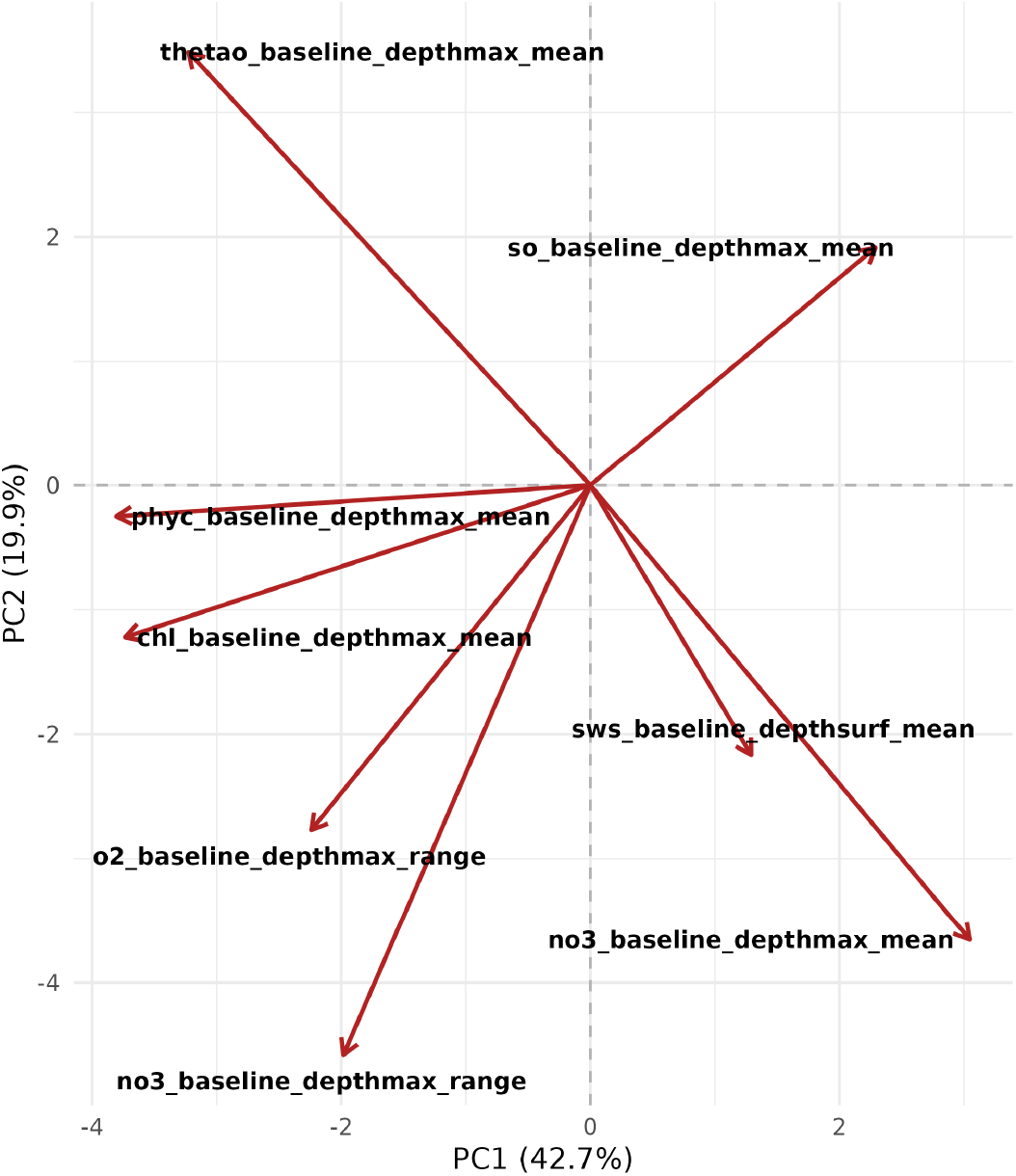
Biplot of the first two Principal Components (PC1 vs PC2), illustrating the environmental space of the study area. Vectors (red arrows) show the direction and magnitude of the original environmental variables. PC1 is strongly driven by SST and PAR, while PC2 aligns with Nitrate and salinity gradients.

In addition to the four climatic PCs, terrain rugosity was included as a static predictor to represent reef structural complexity. It is an important factor for benthic anemone suitability and microhabitat selection Huebner et al. (2012). Bathymetry was used solely for masking (excluding depths > 200 m) to prevent depth artifacts in future projections.

To confirm the statistical independence of the final predictor set (PC1–PC4 + Rugosity), Variance Inflation Factors (VIF) were calculated. All predictors exhibited VIF values < 1.2 (Max VIF = 1.18), far below the threshold of 5 Dormann et al. (2013), confirming suitability for Maximum Entropy modeling.

To assess potential future range shifts, environmental layers were projected into the years 2050 and 2100 under Shared Socioeconomic Pathway (SSP) scenarios SSP1-2.6 (sustainability) and SSP5-8.5 (fossil-fueled development) Riahi et al. (2017). Future climatic variables were projected onto the same PCA axes derived from current conditions for comparability.

### 2.3 Species Distribution Models (SDMs)

We built SDMs for each host sea anemone and anemonefish species using the R package `ENMeval` v2.0 Kass et al. (2021); Muscarella et al. (2014). We utilised the Maxent algorithm implemented via the `maxnet` engine Phillips et al. (2017), suitable for presence-only data, using the log-log (clog-log) output which models occurrence probability.

#### 2.3.1 Data Preparation and Sampling

Prior to modelling, we compiled a binary interaction matrix defining the known associations between the 28 anemone-fish and 10 host anemone species, sourced from the datasets used in previous anemonefish modelling studies Gaboriau et al. (2024). Occurrence records were spatially thinned using a grid-based approach to reduce spatial autocorrelation Boavida et al. (2016), retaining only one unique occurrence per raster grid cell (∼9 km). Species with fewer than 15 spatially unique occurrences after thinning were excluded from further analyses to ensure statistical validity. To characterise the available environment, we generated 10,000 background points (pseudo-absences) for each species. To account for spatial observer bias Phillips et al. (2009), we employed a distance weighted background sampling method. Sampling probability was derived from the geographic and environmental distance to known occurrence records, ensuring that the model differentiates habitat suitability from sampling effort.

#### 2.3.2 Tuning and Evaluation of SDMs

To account for spatial autocorrelation and assess model transferability, we employed a spatial block cross-validation strategy. For each species, the study area was divided into four spatial quadrants defined relative to the median longitude and latitude of the species’ occurrence records Valavi et al. (2019). The models were iteratively trained on three quadrants and evaluated on the fourth to test transferability. Hyperparameter tuning was performed using `ENMeval`. We conducted a grid search testing regularisation multipliers from 0.5 to 4.0 (in steps of 0.5) and four feature class combinations (“Linear”, “Linear-Quadratic”, “Hinge”, and “Linear-Quadratic-Hinge”). The optimal model configuration for each species was selected based on the lowest delta-AICc (Akaike Information Criterion corrected for small sample size) to balance goodness-of-fit with model complexity.

#### 2.3.3 The 3-Stage Biotic Framework

To explicitly test the hypothesis that host availability constrains anemonefish distribution, we implemented a hierarchical, three-stage modelling framework:

1. **Phase 1: Host Anemone Models**. Independent SDMs were built for each of the 10 host sea anemone species using the selected abiotic predictors (PC1–PC4 + Rugosity).
2. **Phase 2: Anemonefish Models**. For each anemonefish species, three competing models were constructed:
  - **Model A (Abiotic-Only):** Modelled using only environmental predictors (PC1–PC4 + Rugosity).
  - **Model B (Biotic-Only):** Modelled using only a species-specific *Biotic Suitability Layer* and Rugosity.
  - **Model C (Combined):** Modelled using abiotic predictors, the *Biotic Suitability Layer*, and Rugosity.

##### Biotic Suitability Layer Construction

For each anemonefish species, we created a specific biotic predictor by calculating the weighted mean suitability of its known host anemone species (derived from the interaction matrix), effectively summarising the landscape of host availability Araújo and Luoto (2007). For future projections, this layer was derived from the *future* projections of host anemones, allowing the model to dynamically capture potential cascade effects where host range shifts constrain fish distribution Schweiger et al. (2010).

#### 2.3.4 Model Uncertainty and Performance

To quantify predictive uncertainty, we bootstrapped the optimal model configuration for each species. We ran 40 iterations for anemonefish and 10 iterations for host anemones, with each iteration using a different random subset of training data. final suitability maps were generated by calculating the ensemble mean and standard deviation across all bootstrap iterations. The predictive performance was assessed using the mean Area Under the Curve (AUC), the True Skill Statistic (TSS), and the Continuous Boyce Index (CBI), a threshold-independent metric particularly robust for presence-only data Hirzel et al. (2006).

### 2.4 Stacking Procedures

To derive community-level richness surfaces for host sea anemones and anemonefish under each scenario and time period, we used a probabilistic stacked SDM approach. We aligned all species-level prediction rasters to a common grid and calculated the cell-wise sum of predicted suitability probabilities across all species. This method provides an estimate of expected species richness (*R*) that retains information on within-cell suitability gradients and avoids the information loss associated with arbitrary binary thresholding Calabrese et al. (2014); Zurell et al. (2020). Changes in community composition were quantified by calculating the delta (Δ) between future and current projections for both individual species and total richness, allowing for the identification of potential range expansions and contractions under future warming scenarios.

#### 2.4.1 Variable Contribution and Driver Analysis

To quantify the mechanistic drivers of distribution, we extracted the regularisation coefficients (Betas) from the fitted MaxNet models. While permutation importance is commonly used, it can be biased by collinearity and spatial autocorrelation Strobl et al. (2008). However, the magnitude of the regularised coefficients provides a direct, mathematically transparent measure of feature importance within the point-process model structure Renner and Warton (2013); Merow et al. (2013). We compared the absolute magnitude of coefficients for environmental gradients (PC1– PC4), Rugosity, and Biotic Suitability to determine if the ecological drivers differed between generalist and specialist guilds.

#### 2.4.2 Spatial Heterogeneity Assessment

To assess the spatial reliability of the predictions, we analysed model performance across the four spatial cross-validation folds. Since the study area was partitioned into distinct longitudinal/latitudinal blocks, consistent performance across all folds indicates high transferability to geographically novel environments Roberts et al. (2017). We aggregated AUC scores by Fold ID for both host and fish models to test the hypothesis that regional uncertainty in host predictions would lead to downstream uncertainty in fish models Wisz et al. (2013).

### 2.5 Statistical Analyses

All post-hoc statistical analyses were performed on the subset of species that successfully passed strict quality control criteria (*N* = 17 fish, *N* =8 hosts). Species were excluded if they failed spatial block cross-validation due to restricted ranges (e.g., *Amphiprion latezonatus*) or possessed insufficient spatially unique occurrence records (*N* < 15) to reliably estimate biotic interactions. Additionally, to ensure the validity of range shift projections, we filtered out any species where the baseline Environmental-Only model failed to achieve a minimum predictive performance of AUC > 0.7 or TSS > 0.4, thresholds commonly cited as the lower bound for useful model accuracy Swets (1988); Allouche et al. (2006). This quality control step ensured that all subsequent analyses of biotic constraints were performed on species for which the fundamental climatic niche could be modelled with a reasonable degree of certainty.

#### 2.5.1 Model Performance and Complexity

To statistically compare the predictive power of the three model formulations (Abiotic-Only, Biotic-Only, Combined), we employed a Linear Mixed Model (LMM) using the `lme4` Bates et al. (2015) and `lmerTest` Kuznetsova et al. (2017) packages. This approach accounts for the nested structure of the data, where multiple performance scores (AUC and TSS) were obtained for each species across bootstrap iterations. We included “Species” as a random intercept to control for inherent variation in modelability among taxa. To test if the inclusion of biotic predictors benefited specific guilds differently, we performed the LMM analysis on the full community and then conducted stratified analyses on the *Generalist* (*N*_*host s*_ ≥ 3) and *Specialist* (*N*_*host s*_ < 3) subsets independently.

#### 2.5.2 Quantifying Biotic Constraints on Range Expansion

To mechanistically explain range shift dynamics (Hypothesis 2), we quantified the biotic constraint on range expansion. For each anemonefish species, we calculated the weighted centroid of its predicted distribution under two conditions:

1. The *Potential Range Shift*, derived from the Abiotic-Only model (representing where the fish could physiologically exist).
2. The *Realised Range Shift*, derived from the Combined model (representing where the fish can exist given host availability).

The difference in poleward displacement between these two centroids represents the magnitude of the *Range Lag* (*km*) caused by the obligate host interaction.

To determine if ecological specialisation exacerbates this constraint, we performed Welch’s two-sample t-tests comparing the mean Range Lag between specialist and generalist species across all future emission scenarios (SSP1-2.6 and SSP5-8.5).

#### 2.5.3 Ghost Habitat Analysis

We quantified the spatial consequence of biotic constraints by identifying areas of Realised Niche Truncation Soberón (2007), termed “Ghost Habitat” here. This is defined as the geographical extent where abiotic conditions are suitable for the symbiont, but the absence of the obligate host prevents occupancy. We converted the continuous probability surfaces from the Abiotic-Only and Combined models into binary presence/absence maps using the Maximum Training Sensitivity plus Specificity (MaxTSS) threshold Liu et al. (2005), averaged across bootstrap iterations. Ghost Habitat was identified as raster cells where:

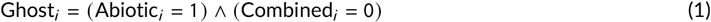

This binary approach provides a conservative estimate of lost colonisation potential, strictly identifying areas where host absence precludes fish establishment despite favourable climate conditions.

#### 2.5.4 Host-fish Spatial Decoupling

To test for the spatial decoupling of the mutualism (Hypothesis 3), we quantified the geographic niche overlap between each anemonefish species and its known hosts under current and future conditions. We used Schoener’s *D* index Warren et al. (2021), a metric ranging from 0 (no overlap) to 1 (identical distribution), calculated using the `ENMeval` package Kass et al. (2021). Change in overlap (Δ*D*) was calculated for every interacting species pair under each future scenario. To assess whether high-emission scenarios destabilise these associations more severely than low-emission scenarios, we analysed the variance in Δ*D* across scenarios, hypothesising that extreme warming (SSP5-8.5) would drive higher heterogeneity and decoupling rates than stabilised scenarios (SSP1-2.6).

#### 2.5.5 Environmental Niche Breadth

To test whether environmental tolerance predicts climate sensitivity, we calculated the realised niche breadth for each anemonefish species. We used Levin’s concentration metrics (*B*_2_), standardised to a scale of 0 to 1, which provides a robust measure of how uniformly a species utilises the available environmental space Levins (1968). *B*_2_ was calculated directly from the continuous suitability scores of the current-day Combined SDM raster for each species:

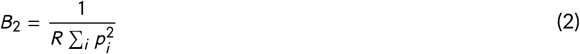

where *p*_*i*_ is the proportional suitability of grid cell *i* relative to the total suitability across the study area, and *R* is the total number of cells. High values (*B*_2_ → 1) indicate broad generalists that utilise a wide range of available habitats, while low values (*B*_2_ → 0) indicate specialists with narrow environmental tolerances. We then performed LMM analysis to determine if current niche breadth (*B*_2_) was a significant predictor of the magnitude of projected range shifts (*km*) under future climate scenarios. We included species identity and emission scenario as random effects to control for repeated measures and the non-independence of projections across different future pathways.

### 2.6 Conservation Prioritisation Framework

To translate ecological predictions into actionable decision-support tools for Marine Spatial Planning (MSP), we developed a prioritisation framework. This analysis turned the species-level probability surfaces into community-level indices to identify climate-resilient refugia and restoration targets. First, we utilised the aggregate species richness surfaces (*R*) derived in Section 2.4 to compute a “Conservation Priority Index” (CPI). The CPI for each grid cell *i* was defined as the product of current biodiversity value and future ecological stability:

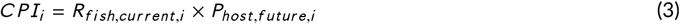

where *R*_*fish*_ is the normalised richness of anemonefish (Combined model) and *P*_*host*_ is the normalised persistence of host anemones under the high-emission scenario (SSP5-8.5, 2100). This index prioritises reefs that currently support diverse fish communities and are projected to retain the biotic infrastructure (hosts) required for their survival, effectively identifying “Climate Refugia” Magris et al. (2018). Second, we calculated a “Restoration Opportunity Index”

(ROI) to use the Ghost Habitat maps. ROI quantifies the gap between the potential climatic niche and the realised biotic niche in the future:

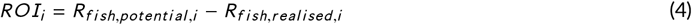

High ROI values indicate areas where the climate is highly suitable for a rich assemblage of fish, but host absence precludes their establishment. These metrics were used to classify the study area into management zones: (1) Refugia (High CPI) for strict protection, and (2) Restoration Frontiers (High ROI) for assisted colonisation.

## 3 RESULTS

The species distribution models developed for this study demonstrated robust predictive capacity, effectively mapping the baseline distributions of both host sea anemones and anemonefish. The results show a clear functional divergence: while environmental variables are the primary drivers of broad-scale distribution for all species, the inclusion of biotic interactions provides an important refinement of the realised niche specifically for specialist species.

### 3.1 Host Sea Anemone Model Performance

SDMs were successfully calibrated for the eight host sea anemone species that met the strict data quality criteria. The abiotic models (PC1–PC4 + Rugosity) achieved moderate-to-high predictive accuracy, with a global mean test AUC of 0.756 (± 0.032 SD) and a mean TSS of 0.454 (± 0.054 SD). Performance varied among species, reflecting differences in niche breadth and record density. The widespread host *Cryptodendrum adhaesivum* achieved the highest predictive performance (Mean AUC = 0.809, TSS = 0.525), suggesting that abiotic predictors effectively capture its realised niche. Conversely, patchily distributed species such as *Heteractis malu* exhibited lower validation scores (Mean AUC = 0.714), indicating that unmeasured local factors, such as specific microhabitat complexity or substrate composition, may limit model precision for these hosts (Table 2).

**TABLE 2.** Predictive performance (mean test AUC ± SD, mean test TSS ± SD) of environmental-only SDMs for host sea anemones.

| Species | Mean Test AUC ± SD | Mean Test TSS ± SD |
| --- | --- | --- |
| *Cryptodendrum adhaesivum* | 0.809 ± 0.012 | 0.525 ± 0.023 |
| *Entacmaea quadricolor* | 0.755 ± 0.021 | 0.416 ± 0.035 |
| *Heteractis aurora* | 0.745 ± 0.015 | 0.401 ± 0.028 |
| *Heteractis crispa* | 0.790 ± 0.018 | 0.541 ± 0.041 |
| *Heteractis magnifica* | 0.767 ± 0.020 | 0.462 ± 0.033 |
| *Heteractis malu* | 0.714 ± 0.033 | 0.455 ± 0.061 |
| *Stichodactyla gigantea* | 0.730 ± 0.025 | 0.425 ± 0.040 |
| *Stichodactyla haddoni* | 0.735 ± 0.019 | 0.407 ± 0.031 |

#### 3.1.1 Drivers of Host Distribution

Analysis of regularisation coefficients revealed that PC1 (representing the primary thermal and light gradient) exhibited the highest mean coefficient magnitude across the host guild (*β* ≈ 1.83). Rugosity also retained explanatory power for hosts (*β* ≈ 1.03) (figure 3).

**FIGURE 3.**
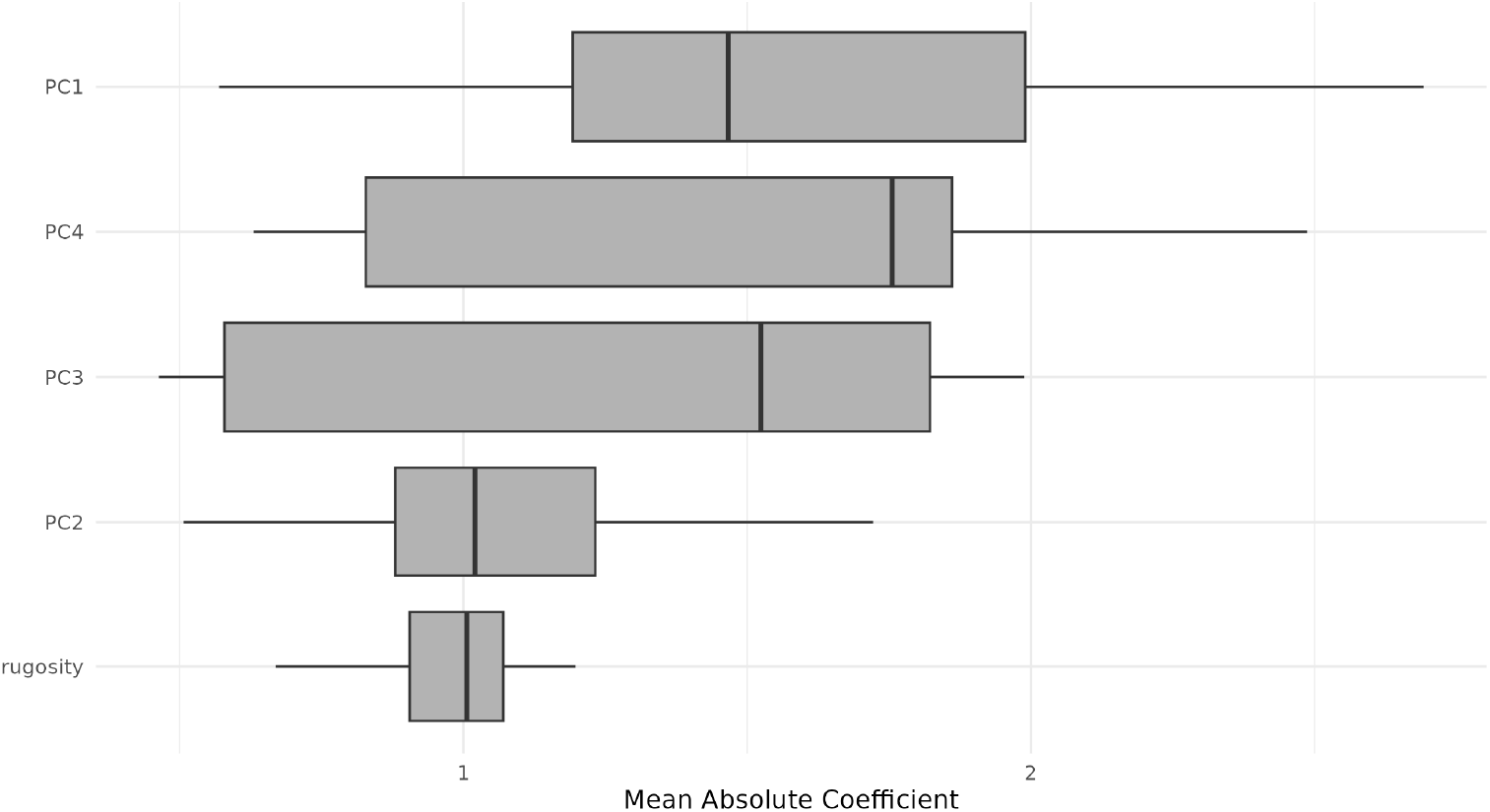
Boxplot for variable Importance for Host Sea Anemone models based on MaxNet coefficients.

**FIGURE 4.**
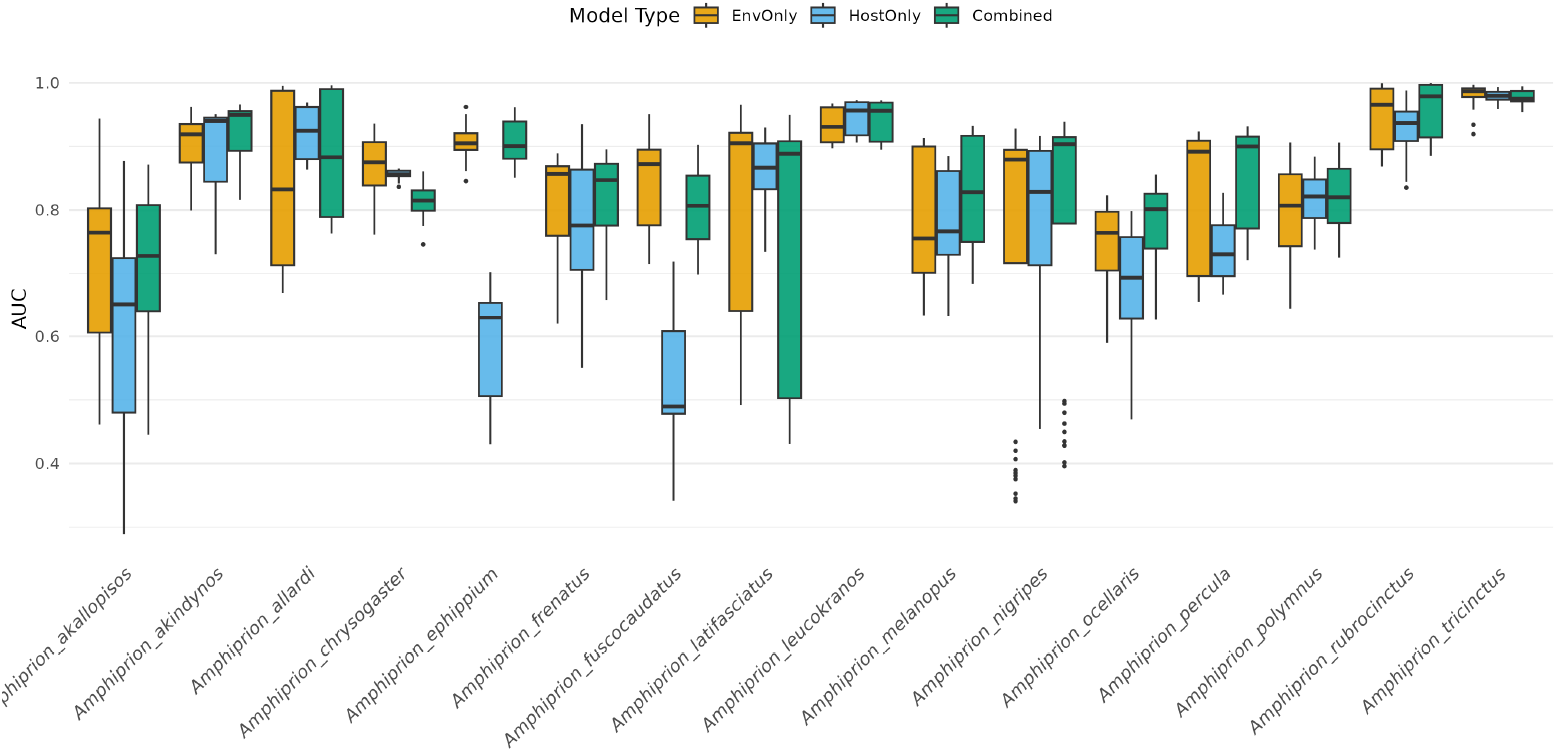
Comparison of Mean AUC for all anemonefish SDMs using Environmental-Only, Biotic (Host) Only, and Combined predictors.

### 3.2 Anemonefish Model Performance

For the anemonefish, we applied a strict quality control filter, excluding any species where the baseline Environmental-Only model failed to achieve a minimum AUC of 0.7. Based on this criterion, *Amphiprion sebae* (AUC = 0.53) was removed from subsequent analyses, leaving a final set of 17 species. Across this final guild, three competing model formulations were evaluated: Environmental-Only (Env-Only), Biotic-Only, and Combined. The Combined models achieved the highest predictive accuracy (Mean AUC = 0.850; Mean TSS = 0.671). Linear Mixed Models (LMM) confirmed that accounting for biotic interactions significantly improved model explanatory power. The Combined model showed a statistically significant increase in both AUC (*β* = +0.01, *t* = 2.16, *p* = 0.031) and TSS (*β* = +0.02, *t* = 3.66, *p* < 0.001) compared to the Environmental-Only baseline. In contrast, models relying solely on host distribution (Host-Only) performed significantly worse than the environmental baseline across all metrics (*β*_*AUC*_ = -0.05, *t* = -11.90, *p* < 0.001; *β*_*TSS*_ = -0.06, *t* = -10.48, *p* < 0.001).

#### 3.2.1 Performance by Ecological Specialisation

When stratified by ecological specialisation, the model improvement was non-uniform. For Generalist species (*n* = 9), the Combined model did not significantly outperform the Env-Only model for either metric (AUC: *p* = 0.534; TSS: *p* = 0.113). In contrast, for Specialist species (*n* = 8), the Combined model significantly outperformed the Env-Only model for both AUC (*β* = +0.022, *t* = 3.28, *p* = 0.001) and TSS (*β* = +0.032, *t* = 3.50, *p* < 0.001). Final predictive performance for the Combined models is detailed in Table 3.

**TABLE 3.** Final predictive performance (mean test AUC ± SD, mean test TSS ± SD) of the Combined SDMs (Abiotic + Biotic predictors) for the 17 analysed anemonefish species.

| Species | Mean Test AUC ± SD | Mean Test TSS ± SD |
| --- | --- | --- |
| *Amphiprion akallopisos* | 0.721 ± 0.109 | 0.446 ± 0.176 |
| *Amphiprion akindynos* | 0.927 ± 0.042 | 0.779 ± 0.114 |
| *Amphiprion allardi* | 0.886 ± 0.101 | 0.759 ± 0.174 |
| *Amphiprion chrysogaster* | 0.815 ± 0.024 | 0.676 ± 0.030 |
| *Amphiprion chrysopterus* | 0.927 ± 0.014 | 0.748 ± 0.032 |
| *Amphiprion ephippium* | 0.908 ± 0.033 | 0.770 ± 0.085 |
| *Amphiprion frenatus* | 0.815 ± 0.077 | 0.554 ± 0.139 |
| *Amphiprion fuscocaudatus* | 0.803 ± 0.061 | 0.607 ± 0.138 |
| *Amphiprion latifasciatus* | 0.765 ± 0.201 | 0.642 ± 0.242 |
| *Amphiprion leucokranos* | 0.942 ± 0.029 | 0.861 ± 0.079 |
| *Amphiprion melanopus* | 0.829 ± 0.080 | 0.582 ± 0.157 |
| *Amphiprion nigripes* | 0.794 ± 0.202 | 0.558 ± 0.242 |
| *Amphiprion ocellaris* | 0.835 ± 0.090 | 0.623 ± 0.151 |
| *Amphiprion percula* | 0.914 ± 0.015 | 0.715 ± 0.029 |
| *Amphiprion polymnus* | 0.911 ± 0.020 | 0.733 ± 0.050 |
| *Amphiprion rubrocinctus* | 0.977 ± 0.011 | 0.918 ± 0.023 |
| *Amphiprion tricinctus* | 0.965 ± 0.023 | 0.879 ± 0.051 |

### 3.3 Variable Contribution by Guild

Analysis of MaxNet regularisation coefficients indicated distinct differences in predictor importance between guilds (figure 5). For Generalist species, the Biotic Suitability term was the dominant predictor (Median *β* ≈ 3.1), exceeding the magnitude of environmental variables. In contrast, Specialist species showed high coefficient values for both PC1 (Climate, Median *β* ≈ 3.8) and Biotic Suitability (Median *β* ≈ 1.7). While Biotic Suitability remained a strong predictor, the absolute contribution of the primary thermal gradient (PC1) was consistently higher for specialists than for generalists.

**FIGURE 5.**
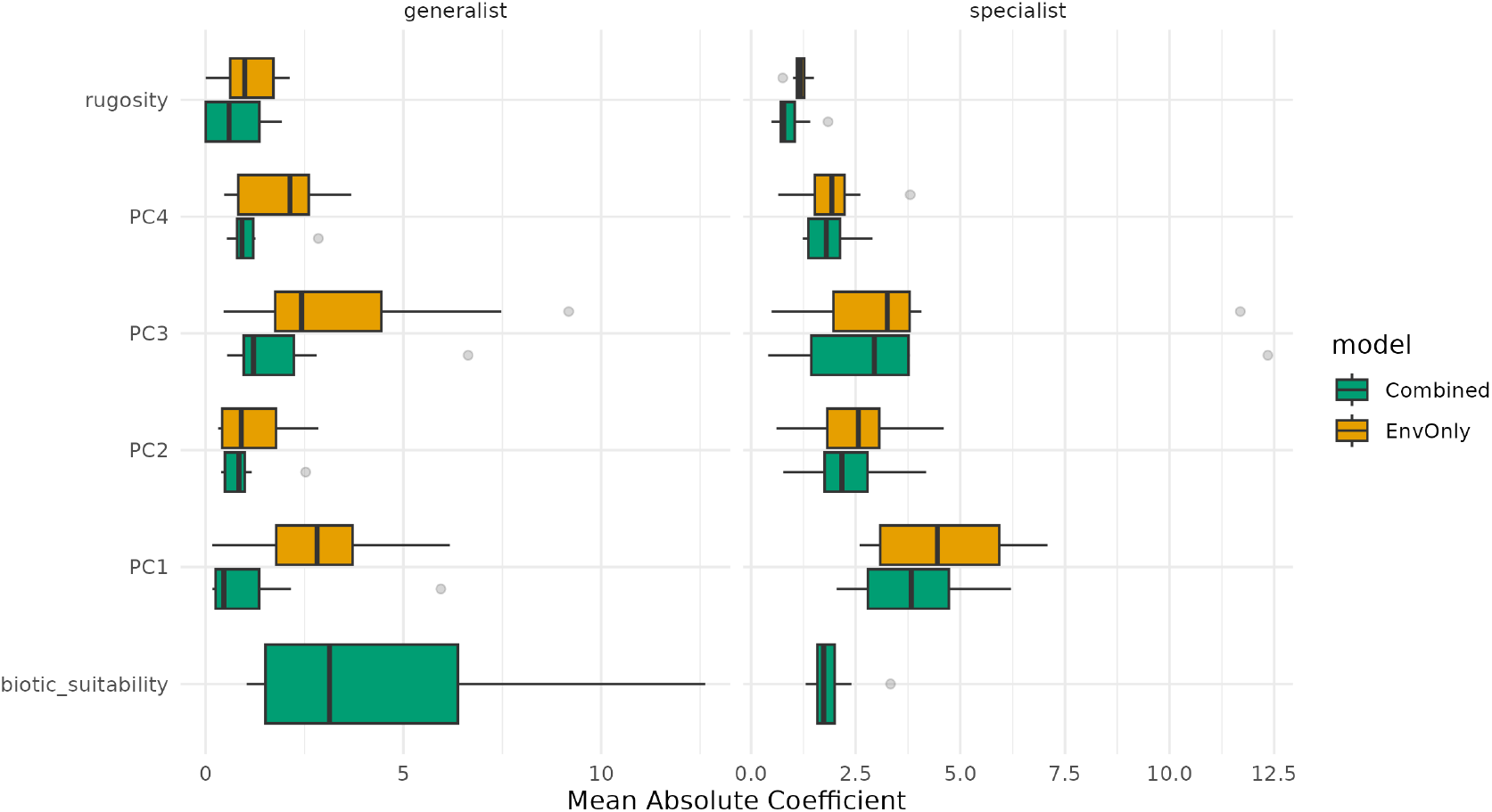
MaxNet coefficient magnitudes for Anemonefish models. Left (Generalists), Right (Specialists)

**FIGURE 6.**
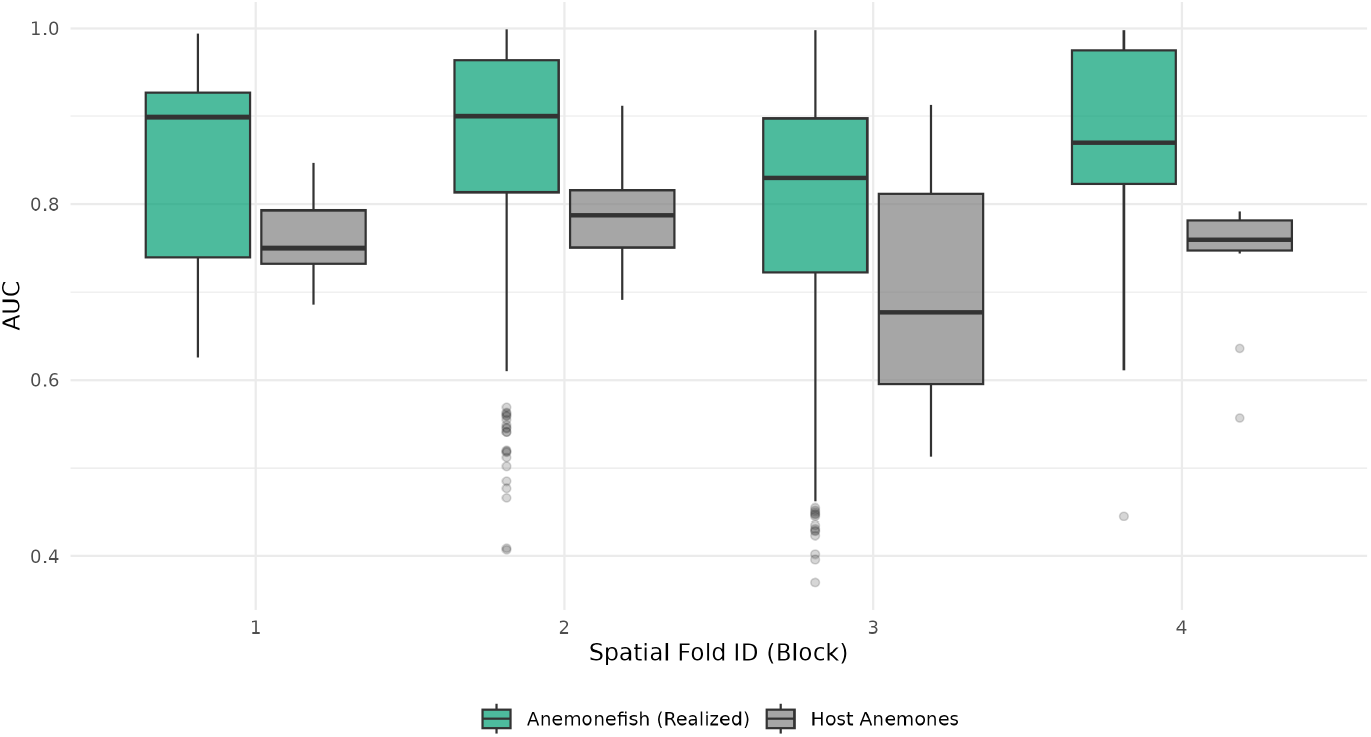
Distribution of AUC scores across spatial folds. Both Host and fish models exhibit a dip in performance in Fold 3.

### 3.4 Spatial Heterogeneity and Error Propagation

To assess spatial reliability, model performance was analysed across four spatial quadrants based on the median coordinates of occurrence records. This analysis revealed spatial variation in predictive accuracy. For 50% of the analysed species (12/24), Spatial Fold 3 (representing the South-East quadrant relative to the species’ range center) yielded the lowest mean AUC (0.776 for fish). This pattern was consistent across both the anemonefish (*N* = 16, 8 worst in Fold 3) and Host Anemone (*N* = 8, 4 worst in Fold 3) guilds, where Host models also exhibited their lowest mean accuracy in this fold (Mean AUC ≈ 0.708).

### 3.5 Biotic Constraints on Range Expansion

To mechanistically explain the divergence between potential (climate-only) and realised (host-constrained) range shifts, we quantified the Biotic Constraint as the difference in centroid displacement between the Environmental-Only and Combined models. The results indicate that obligate host interactions act as a substantial but highly variable constraint on climate-driven range expansion. Under the high-emission scenario (SSP5-8.5, 2100), the mean biotic constraint (or “Range Lag”) for specialist species was 97.4 km (± 305.0 SD), compared to 35.2 km (± 982.1 SD) for generalist species. This suggests that while specialists generally face a constraint, the response among generalists is extremely diverse, with some species exhibiting severe lags while others track climate velocity effectively. Individual responses highlighted this variability. The specialist *Amphiprion akallopisos* exhibited a biotic lag of 212.4 km under SSP5-8.5 (2100), losing significant potential range due to host immobility. In contrast, the generalist *Amphiprion akindynos* showed a negative constraint (-139.5 km), implying that its host interactions actually facilitated range expansion into marginal high-latitude habitats (figure 7). Despite the trend of higher mean lag in specialists, Welch’s two-sample t-tests revealed no statistically significant difference in the magnitude of biotic constraint between the two guilds (*t* = 0.17, *df* = 8.8, *p* = 0.87; Table 4). This lack of significance is driven by the extreme variance within the generalist guild, indicating that while biotic constraints are pervasive, their magnitude is determined more by species-specific host associations than by broad ecological specialisation categories.

**TABLE 4.** Statistical comparison (Welch’s t-test) of the mean Biotic Constraint (km) between specialist and generalist anemonefish across four future climate scenarios.

| Scenario | Mean Lag (Gen) | Mean Lag (Spec) | t-value | df | p-value |
| --- | --- | --- | --- | --- | --- |
| SSP1-1.9 (2050) | 10.8 km | 64.6 km | -0.37 | 9.19 | 0.72 |
| SSP1-1.9 (2100) | 0.7 km | 51.7 km | -0.42 | 9.18 | 0.69 |
| SSP5-8.5 (2050) | 11.8 km | 55.1 km | -0.36 | 9.34 | 0.73 |
| SSP5-8.5 (2100) | 35.2 km | 97.4 km | 0.17 | 8.82 | 0.87 |

**FIGURE 7.**
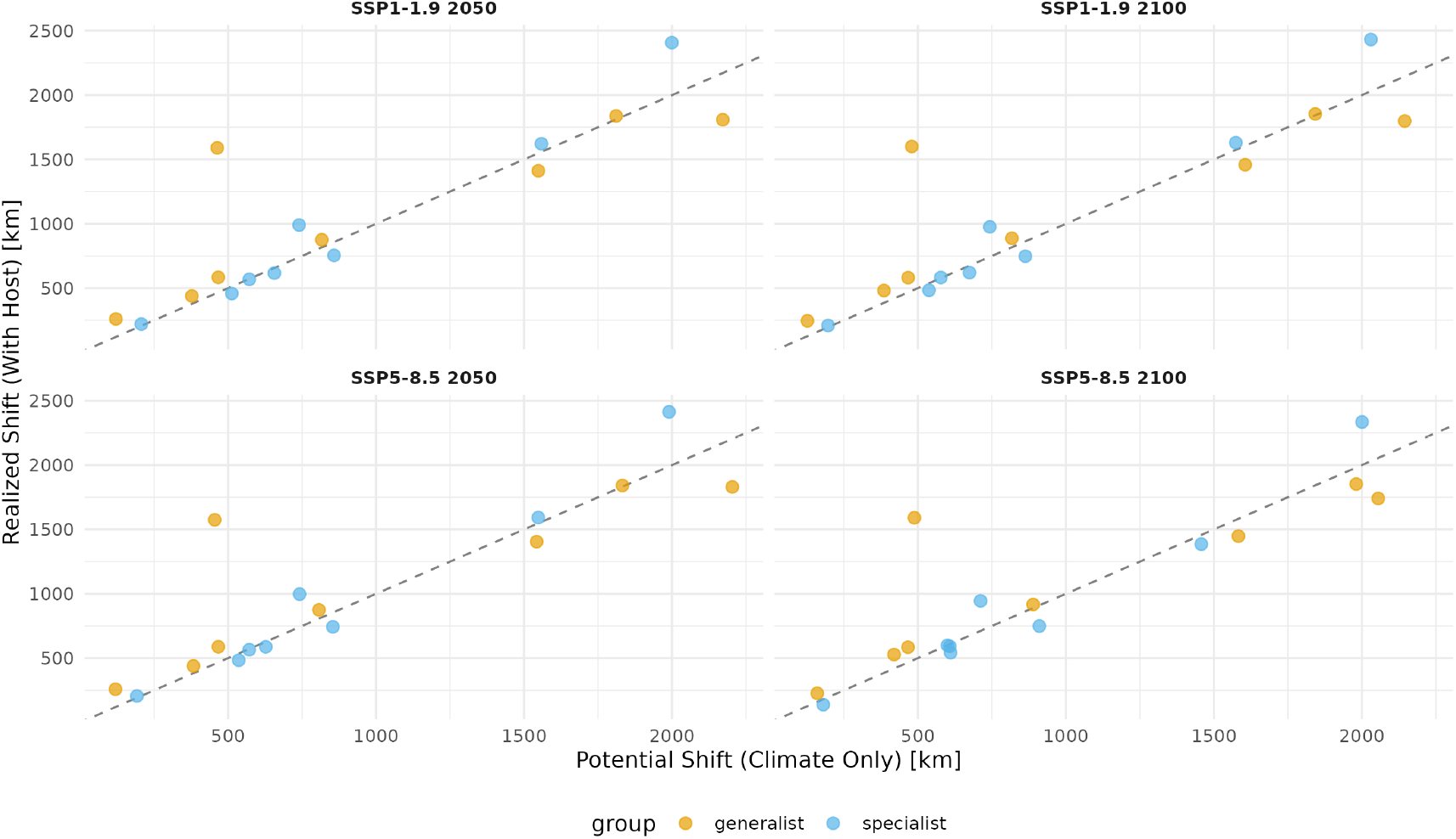
The Biotic Constraint on Range Expansion (SSP5-8.5, 2100). The dashed 1:1 line represents the null hypothesis where host presence imposes no constraint (Realised Shift = Potential Shift). Points below this line indicate species that are lagging behind their climatic potential due to host dependence. The vertical distance from the line represents the magnitude of this biotic lag.

### 3.6 Ghost Habitat and Niche Decoupling

While the geometric range lag was variable, the spatial consequence of host dependence was unambiguous. The primary mechanism driving range loss is the formation of Ghost Habitats. Our analysis shows imbalance in the burden of Ghost Habitat. Specialist species are projected to face a persistently high deficit of usable space, with an aggregate Ghost Habitat area of approximately 3.2 million km^2^ across all future scenarios. In contrast, generalist species face a significantly lower burden of approximately 0.7 million km^2^ (figure 8). The projected changes in total suitable habitat area by 2100 under SSP5-8.5 reveal a guild of clear winners and losers (Table 5). While some generalists like *Amphiprion allardi* are projected to expand their range by over 34%, the majority of specialists face severe contractions. *Amphiprion ocellaris* (-54.2%) and *Amphiprion frenatus* (-51.6%) are projected to lose over half of their currently suitable habitat.

**TABLE 5.** Projected changes in suitable habitat area (km^2^) for all anemonefish species under the high-emission scenario (SSP5-8.5) by the year 2100. Negative values indicate range contraction.

| Species | Guild | Current Area (km<sup>2</sup>) | Change (%) |
| --- | --- | --- | --- |
| *Amphiprion akallopisos* | Specialist | 2,903,457 | -20.8% |
| *Amphiprion akindynos* | Generalist | 1,255,112 | -9.2% |
| *Amphiprion allardi* | Generalist | 620,409 | +34.5% |
| *Amphiprion chrysogaster* | Generalist | 660,580 | -59.6% |
| *Amphiprion ephippium* | Specialist | 2,803,870 | -47.4% |
| *Amphiprion frenatus* | Specialist | 3,664,489 | -51.6% |
| *Amphiprion fuscocaudatus* | Generalist | 3,670,783 | -29.3% |
| *Amphiprion latifasciatus* | Generalist | 1,977,099 | +22.6% |
| *Amphiprion leucokranos* | Generalist | 2,126,341 | -63.4% |
| *Amphiprion melanopus* | Specialist | 2,777,648 | -39.4% |
| *Amphiprion nigripes* | Specialist | 1,947,768 | -21.7% |
| *Amphiprion ocellaris* | Specialist | 3,919,688 | -54.2% |
| *Amphiprion percula* | Specialist | 2,408,026 | -50.8% |
| *Amphiprion polymnus* | Generalist | 2,754,056 | -27.3% |
| *Amphiprion rubrocinctus* | Specialist | 652,074 | -51.0% |
| *Amphiprion tricinctus* | Generalist | 948,130 | -63.2% |

**FIGURE 8.**
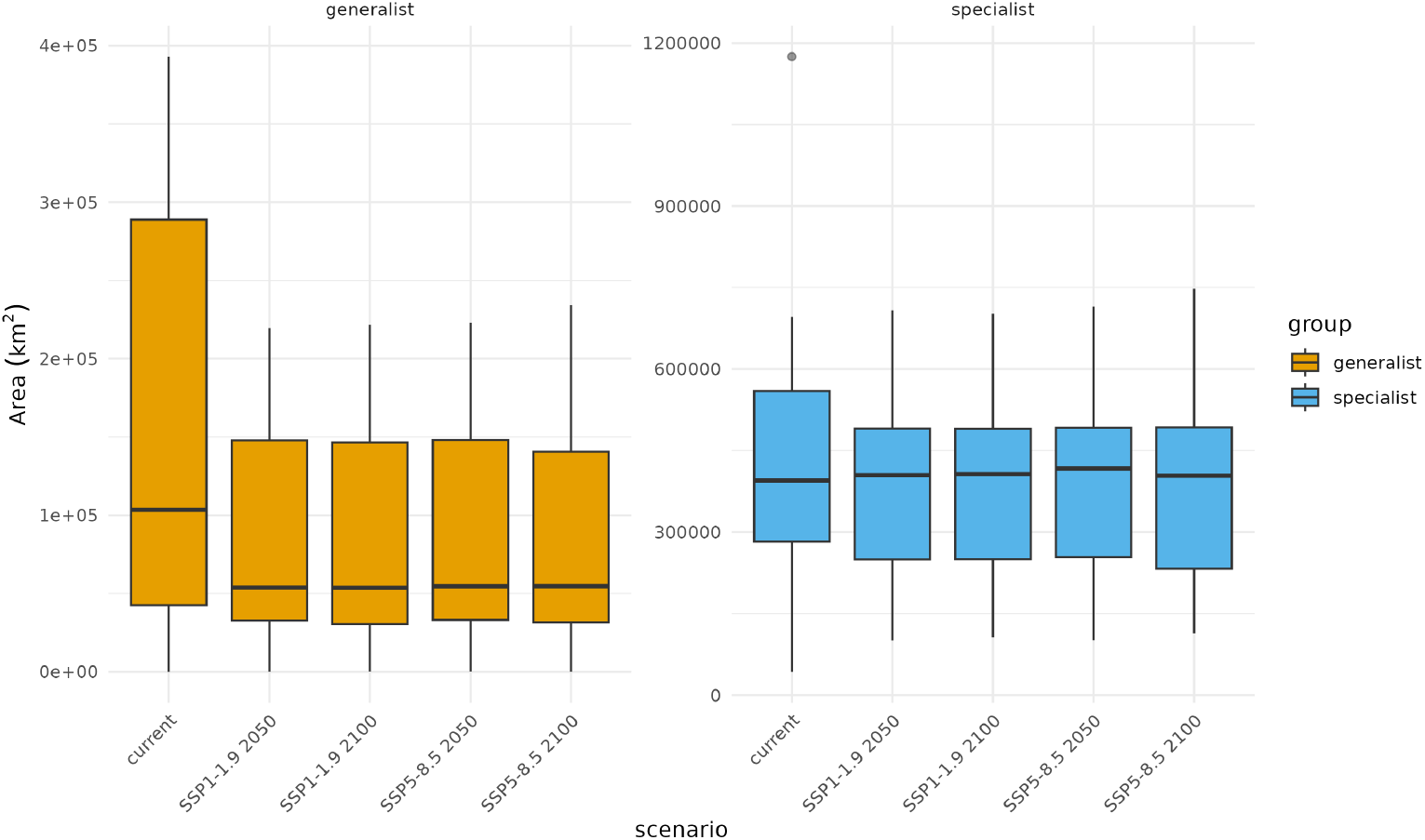
Trends in total Ghost Habitat area (km^2^) across emission scenarios. Specialists (blue) face a consistently higher burden of inaccessible habitat compared to generalists (orange).

**FIGURE 9.**
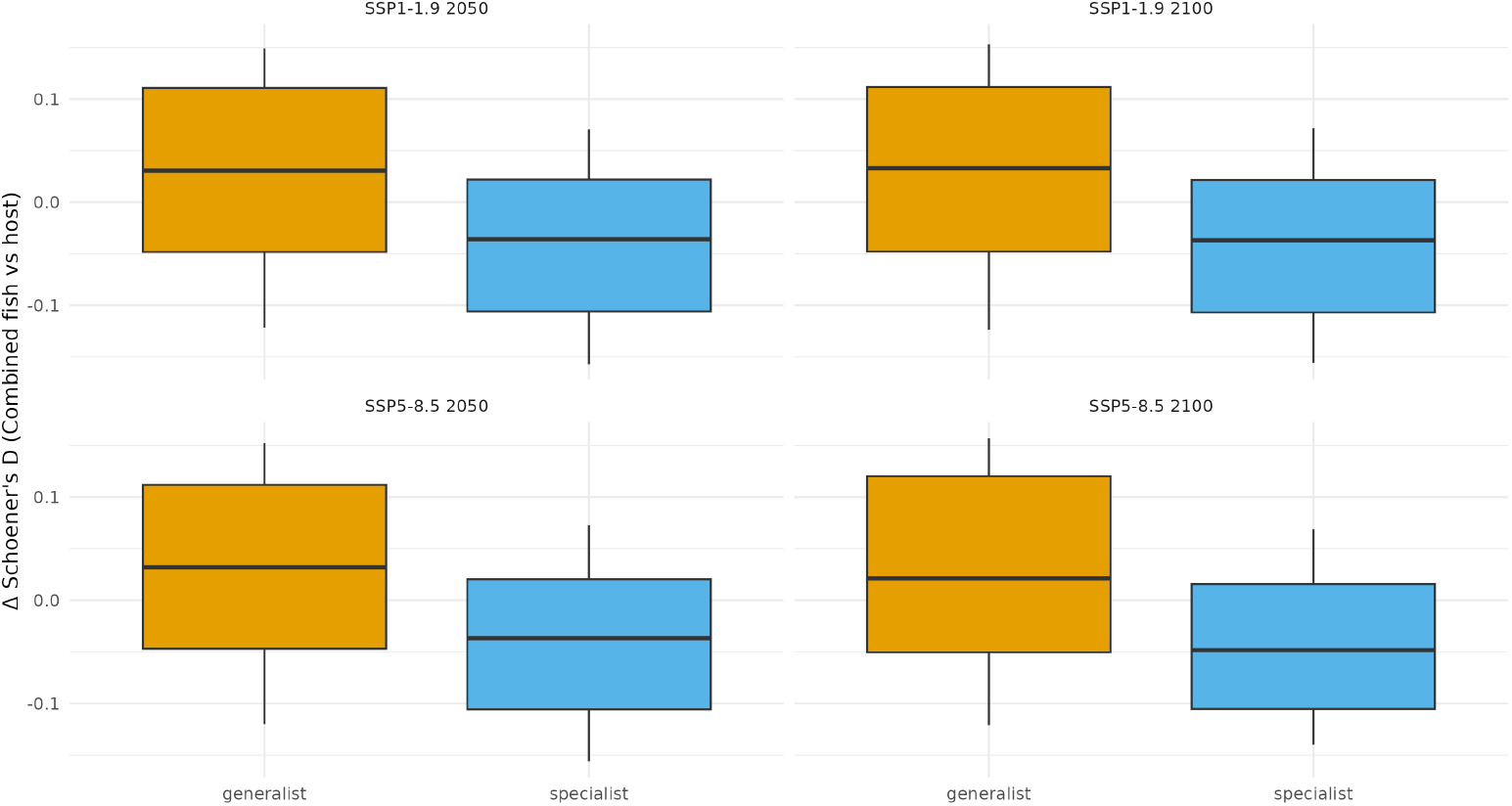
Change in realised niche overlap (Δ*D*) between anemonefish and their hosts under future climate scenarios.

This loss of realisable area is mirrored by a spatial decoupling of the mutualism. Analysis of niche overlap change (Δ*D*) reveals divergent trajectories. Under SSP5-8.5 (2100), generalist species are projected to maintain stable spatial overlap with hosts (Mean Δ*D*_*comb*_ ≈ +0.025). Conversely, specialist species are projected to suffer a consistent decline in overlap (Δ*D*_*comb*_ ≈ −0.044). This negative trend signifies a progressive spatial mismatch, where the climatic envelopes of specialists and their specific hosts are drifting apart.

### 3.7 Environmental Niche Breadth and Climate Sensitivity

finally, we investigated whether a species’ intrinsic environmental tolerance predicted its sensitivity to future climate change. We calculated Levin’s niche breadth (*B*_2_) for all species based on their realised (Combined) niche. The LMM revealed a marginally significant negative correlation between niche breadth and the magnitude of projected poleward range shift (*p* = 0.073, *β* = -2130.8). Species with narrower environmental niches (*low B*_2_) were projected to experience substantially larger latitudinal shifts in suitability than broad-niche generalists. For instance, the range-restricted specialist *Amphiprion nigripes* (*B*_2_ = 0.30) is projected to shift over 2,800 km, whereas widespread generalists typically exhibited shifts of less than 1,000 km. This inverse relationship suggests that species with narrower thermal and chemical tolerances are subject to higher effective climate velocity, requiring greater geographical displacement to remain within their realised niche limits (figure 10).

### 3.8 Community Richness and Conservation Zoning

The stacking of individual SDMs revealed distinct spatial patterns in community vulnerability. Under current conditions, species richness for both guilds peaks in the Coral Triangle ecoregion. However, future projections under SSP5-8.5 (2100) indicate a divergence: host richness is projected to contract severely across the equatorial tropics, while fish climatic suitability remains widespread. The “Conservation Priority Index” (CPI) map identifies a fragmented network of resilient reefs where high current biodiversity intersects with high future host persistence. These refugia are primarily restricted to the subtropical bands, specifically the Kuroshio Current expansion zone and the Southern Great Barrier Reef (figure 11A). Conversely, the “Restoration Opportunity Index” (ROI) reveals that Ghost Habitat is prevalent across the equatorial Indo-Pacific (figure 11B). High ROI values dominate the tropical core, identifying vast areas where the climate remains suitable for up to 80% of the anemonefish guild, yet host collapse renders the habitat functionally sterile. The final Climate-Ready Zoning Map (figure 11C) shows these metrics more clearly. It classifies approximately 15% of the study area as high-priority Refugia warranting strict protection, while highlighting extensive Restoration Frontiers throughout the equatorial belt where active intervention (e.g., host transplantation) is required to create the latent climatic potential.

**FIGURE 10.**
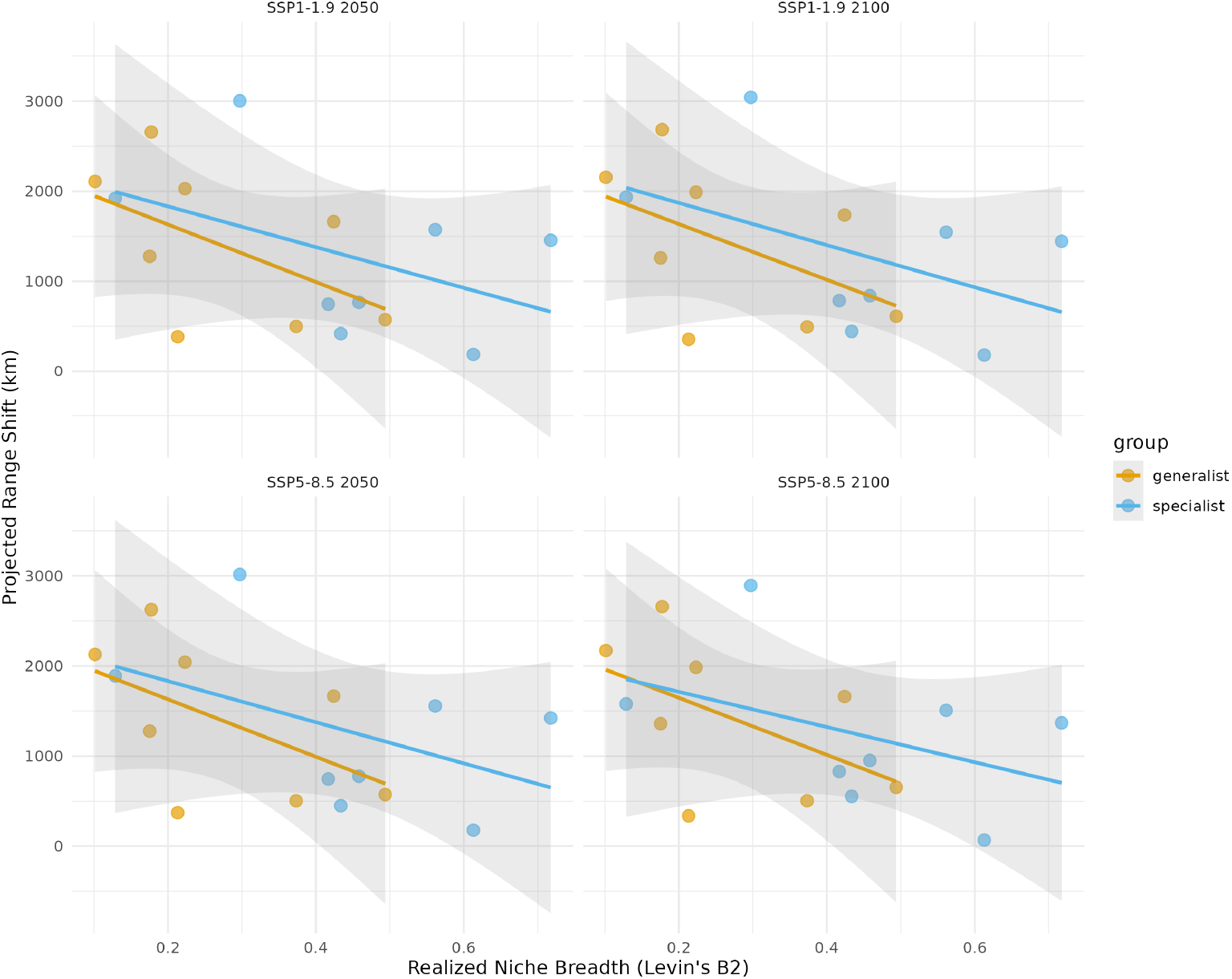
Relationship between current realised niche breadth (Levin’s *B*_2_) and the magnitude of projected poleward range shift (*km*) under future climate scenarios. The negative trend indicates that species with narrower realised niches (specialists) are generally projected to shift further poleward than broad-niche generalists.

**FIGURE 11.**
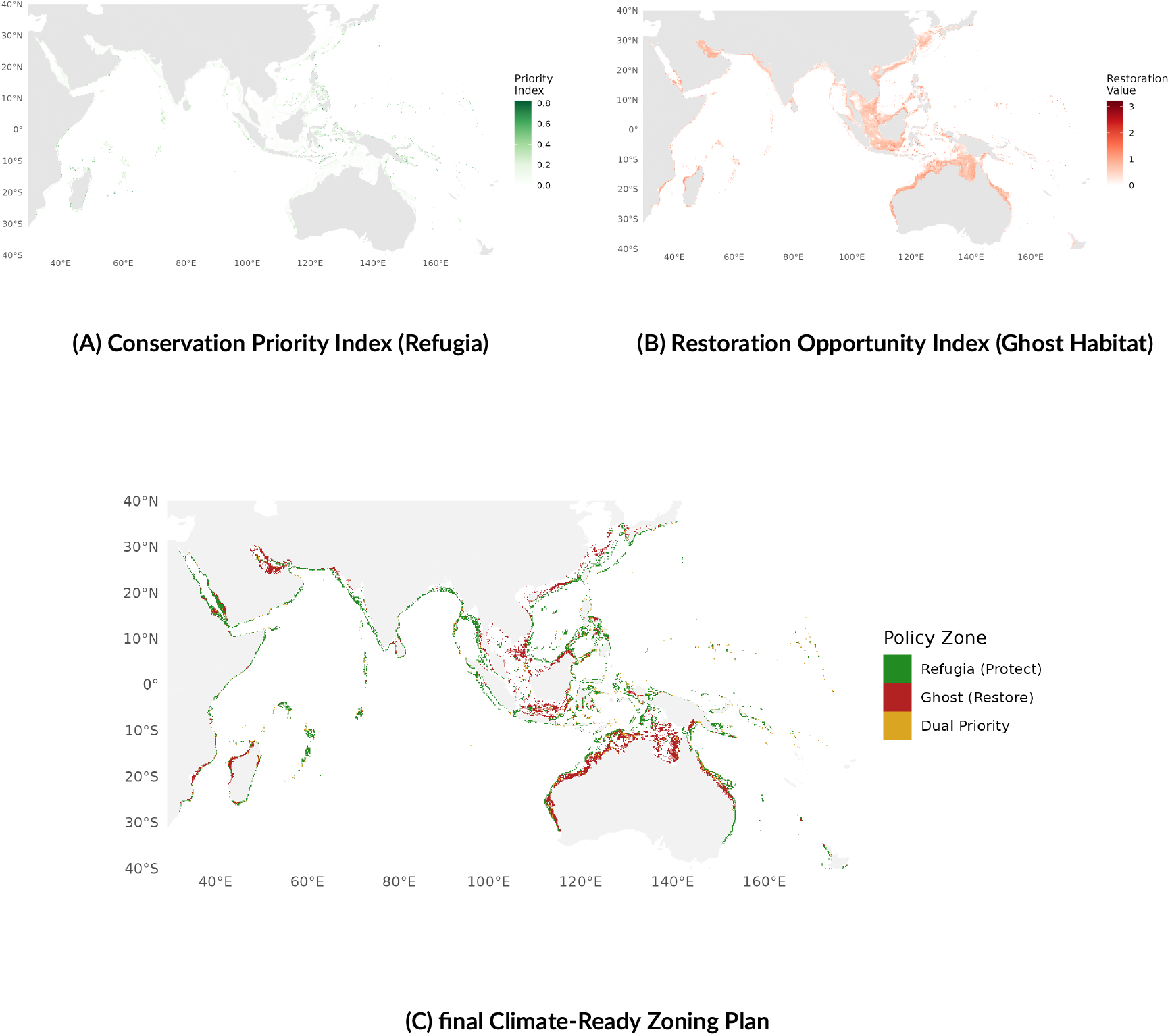
Climate-Ready Conservation Zoning Proposal derived from SSP5-8.5 projections. (A) Refugia (Green): Areas of high stability, primarily in subtropical latitudes. (B) Restoration Frontiers (Red): Areas of high climatic suitability but low host availability. (C) Zoning Synthesis: Prioritising protection in the north/south refugia and intervention in the equatorial restoration zones.

## 4 DISCUSSION

Our main finding is that biotic interactions are not secondary modifiers of local abundance but are fundamental determinants of species distributions at the macro-scale. By integrating host sea anemone distributions into SDMs for 17 anemonefish species, we demonstrated that host availability acts as an important biotic filter that overrides broad-scale environmental suitability for many species. These results challenge the Eltonian Noise Hypothesis, which posits that biotic interactions effectively average out at large spatial scales, leaving climate as the sole driver of distribution Soberón and Nakamura (2009). Instead, our findings support a hierarchical interpretation where the abiotic environment structures host distributions, and host availability in turn strictly defines the realised niche of the fish symbionts Soberon and Peterson (2005a). The inclusion of biotic data improved the predictive accuracy of the models, but this benefit was strongly asymmetric. Specialist species showed significant performance gains, suggesting that climate-only models drastically over-predict their potential range. However, generalist species exhibited negligible improvement, suggesting their distributions are largely resolved by climate alone. This divergence was reflected in the spatial projections for future climate scenarios. Obligate host interactions acted as a biotic constraint, creating substantial lags between potential climatic shifts and realisable range expansions. However, the manifestation of this constraint was complex. While the geometric range lag was highly variable across species, the loss of usable habitat was unambiguous: specialists are projected to lose access to vast areas of climatically suitable ocean (Ghost Habitat) simply due to the immobility of their hosts. Furthermore, we identified a link between environmental tolerance and climate sensitivity. Species with the narrowest realised niches are projected to experience the highest effective climate velocities, requiring geographically implausible dispersal distances to track their fleeing optima. Collectively, these findings suggest that the breakdown of mutualisms, driven by the spatial decoupling of fish and host climatic envelopes, is a greater threat to anemonefish persistence than physiological thermal limits alone.

### 4.1 Biotic Interactions as Macro-Ecological Drivers

The Eltonian Noise Hypothesis states that biotic interactions are transient or local, leaving climate as the sole driver of distribution at continental scales Soberón and Nakamura (2009). Our results challenge this view for obligate mutualists, but with nuance: the scale of the biotic signal depends on the degree of specialisation. The significant improvement in model performance (AUC/TSS) for specialist anemonefish when host data were included confirms that their distributions are not fully resolved by abiotic variables. For species like *Amphiprion nigripes* or *A. rubrocinctus*, climate suitability alone overpredicted their range. Our coefficient analysis supports this mechanism: specialists exhibited high sensitivity to both PC1 (thermal gradients) and biotic suitability, indicating they are constrained by a dual filter of physiological limits and host availability. This overprediction error in abiotic-only models shows that the realised niche of specialists is a strict subset of the fundamental climatic niche, created by the biotic interaction Soberon and Peterson (2005b). In contrast, the lack of significant improvement for generalist species supports a partial validity of the Eltonian framework for broad-niche taxa. However, variable importance analysis revealed that biotic suitability was actually the dominant predictor for generalists, often exceeding environmental variables. This apparent contradiction (high variable importance but low model improvement) suggests extreme collinearity. Generalists such as *Amphiprion clarkii* or *A. akindynos* associate with up to 10 different host species Fautin and Allen (1997). This functional redundancy effectively smooths out the biotic constraint; the aggregate distribution of the host assemblage converges with the broad environmental gradients of the reef itself, making the biotic predictor redundant to climate variables. This finding resolves conflicting debates in an SDM literature Wisz et al. (2013) by demonstrating that the necessity of biotic predictors is not a binary choice but a function of ecological specialisation.

### 4.2 Biotic Constraints and Ghost Habitats

A central hypothesis of this study was that the limited dispersal capacity of benthic hosts would constrain the pole-ward range expansion of their mobile fish symbionts. Our analysis confirms the existence of this Biotic Constraint, but suggests that its manifestation is more complex than a simple binary between specialists and generalists. The mechanistic basis for this constraint lies in the fundamental life-history divergence between the partners, creating a dispersal asymmetry Schweiger et al. (2010). Anemonefish possess a pelagic larval phase capable of long-distance dispersal (10–100 km per generation) Pinsky et al. (2010), theoretically allowing them to track rapid climate velocities. In contrast, host sea anemones are sessile benthic invertebrates restricted by the physiological limits of their endosymbiotic algae. As sea surface temperatures rise, hosts face a depth-light trap; they cannot retreat to cooler deep waters without losing essential solar irradiance, nor can they migrate poleward rapidly enough to keep pace with the shifting isotherms Fautin and Allen (1997). Our coefficient analysis validates this physiological bottleneck, identifying PC1 (Temperature/Light) as the dominant driver of host distribution (*β* ≈ 1.83).

Quantitatively, this resulted in the disparity between potential and realised range shifts (Range Lag). However, this geometric lag did not differ significantly between specialists and generalists (*p* > 0.05) due to extreme variance within the generalist guild. This variance likely stems from the idiosyncratic biogeography of the host species themselves. Some generalists associate with temperate-tolerant hosts (e.g., *Entacmaea quadricolor*) that are already establishing high-latitude footholds Yamano et al. (2011), facilitating the fish’s expansion. Others are tethered to tropical-restricted hosts, creating large lags. This suggests that generalism is not a universal buffer against climate change; a generalist is only as resilient as its most poleward-tolerant host Schweiger et al. (2010). While, the geometric shift was variable, the spatial consequence was clear. The Ghost Habitat analysis shows the strong evidence of biotic vulnerability. Specialists are projected to lose access to over 3.2 million km^2^ of climatically suitable ocean by 2100 simply because their hosts cannot migrate fast enough. This phenomenon creates regions that appear perfect in climate-only models but remain biologically empty. For species like *Amphiprion akallopisos*, this represents a functional reduction of their potential future range by nearly 21%, a loss which is invisible to traditional assessments. This highlights that for obligate mutualists, the greatest threat is not necessarily physiological stress, but the spatial decoupling of the interaction itself, leading to a silent extinction of potential range Schweiger et al. (2010).

### 4.3 Intrinsic Tolerance and Vulnerability to Niche Decoupling

Beyond the external constraints imposed by host immobility, our results identify a second vulnerability for specialist species; their narrow environmental tolerance. This is mechanistically supported by the variable importance analysis, which showed that specialists exhibit large sensitivity to the primary climate gradient (PC1 Median *β* ≈ 3.8), confirming they are physiologically confined to a narrow thermal window. We found a significant negative relationship between realised niche breadth (*B*_2_) and the magnitude of the projected range shift. Specialists with narrow thermal and chemical tolerances (e.g., *Amphiprion nigripes, B*_2_ = 0.30) are projected to undergo latitudinal displacements exceeding 2,800 km to track their climatic optima, whereas broad-niche generalists (e.g., *Amphiprion sebae, B*_2_ = 0.81) often require shifts of less than 500 km.This disparity is best understood through Climate Velocity; the speed at which a specific set of environmental conditions moves across the landscape Burrows et al. (2011). Species with narrow niches are effectively tracking a very thin slice of environmental space. As the ocean warms, these narrow climatic bands migrate rapidly poleward, often traversing vast geographic distances to remain constant. In contrast, generalists possess a broad physiological buffer. Their wider tolerance allows them to persist *in situ* or undergo only minor distributional adjustments, as the future conditions at their current locations often remain within their fundamental envelope.

This creates a vulnerability to niche decoupling for range-restricted anemonefish. They are subject to the highest effective climate velocities (requiring rapid, long-distance migration to remain within their physiological limits) while simultaneously being locked to the most dispersal-limited hosts. The requirement for an anemonefish species like *A. nigripes* to shift nearly 3,000 km in less than a century is biologically implausible for coral reef species with larval durations of 10–20 days Pinsky et al. (2010). The widening gap between the required dispersal for survival and the realised dispersal potential suggests that for many specialists, range expansion is not a viable adaptation strategy Devictor et al. (2012). Instead, their persistence will depend entirely on their ability to adapt locally to conditions outside their current realised niche, a capacity that their historically narrow tolerance suggests is limited. Furthermore, this finding has evolutionary implications. The correlation suggests that specialisation itself acts as an evolutionary dead-end during periods of rapid climate change. While narrow niches allow for efficient exploitation of stable environments, they strip the species of the flexibility required to navigate the current and future accelerated environmental change. This supports the theoretical prediction that generalists will be the winners of the current extinction crisis, inheriting the reefs left vacant by the specialist losers Clavel et al. (2011).

### 4.4 Spatial Uncertainty and Environmental Complexity

The reliability of species distribution projections is not spatially uniform. Our analysis of spatial cross-validation folds showed a consistent bias: predictive accuracy was lowest in Spatial Fold 3 (representing the North-West quadrant relative to the species’ range center) for 50% of analysed species. This region, often corresponding to the semi-enclosed basins and complex archipelagos of the Indo-Pacific (e.g., South China Sea, Indonesian Throughflow), shows a challenge for broad-scale models, likely due to spatial non-stationarity in environmental drivers Miller and Hanham (2011). The low performance of both host and fish models in this quadrant suggests potential compounding uncertainty: the difficulty in mapping patchy benthic hosts in these heterogeneous environments propagates downstream, limiting the accuracy of fish predictions Wisz et al. (2013). This finding shows that for conservation planning SDMs may be least reliable in regions of high environmental dynamism, requiring improved precautionary buffers when designating protected areas in these zones Kujala et al. (2013).

### 4.5 Conservation Implications: A Host-first Strategy

The identification of the Biotic Constraint and Ghost Habitat redirects the conservation focus for the anemonefish-anemone guild. Traditional management strategies that focus on climate-proofing reefs based solely on fish physiology, such as protecting cool-water refugia identified by thermal stress anomalies, may be insufficient if those refugia lack the specific hosts required for colonisation. Our results suggest that the fish are not the primary victims of thermal stress; the hosts are. Host sea anemones are prone to bleaching and mortality under thermal stress Hobbs et al. (2013), reducing habitat quality even where fish could otherwise persist. Therefore, conservation interventions should prioritise a host-first strategy, protecting the persistence of host populations, particularly at the leading edges of their range where expansion is most important to releasing the constraint on fish redistribution. A practical strength of this study is the translation of abstract biotic constraints into spatially explicit management tools. The Host Limited Niche maps (Ghost Habitat) identify specific reefs that are already climatically suitable for anemonefish but remain ecologically inaccessible due to host absence (figure 8). These locations should be treated as priority targets for: (i) intensified field validation to confirm host absence, and (ii) host-focused management actions, such as minimising local stressors (e.g. sedimentation, physical damage) that reduce host survivorship Brown et al. (2013).Furthermore, the projected vulnerability to niche decoupling for specialist species raises the potentially controversial discussion of Assisted Colonisation. For species like *Amphiprion nigripes*, which faces a dispersal gap of thousands of kilometres, natural range expansion is unlikely to keep pace with warming. The identification of Host Limited Niche hotspots provides a roadmap for experimental translocations. If hosts can be successfully propagated and restored on these climatically suitable but empty reefs, the dependent fish would likely follow, effectively releasing the constraint on their adaptation. However, such interventions carry non-trivial ecological risks and should be approached with risk-benefit framework, starting with small-scale pilot studies on restoration-ready reefs Hoegh-Guldberg et al. (2008).

The spatial prioritisation analysis (figure 11) provides a concrete roadmap for implementing a host-first strategy. The identification of stable refugia in the subtropical expansion zones confirms the importance of the Southern Great Barrier Reef and Kuroshio Current as major climate corridors. These areas, having high CPI scores, should be the primary targets for Marine Protected Area (MPA) expansion to safeguard the reproductive sources of the future Beger et al. (2010). Simultaneously, the Restoration Opportunity Index (ROI) identifies Restoration Frontiers across the equatorial tropics. This metric shifts the narrative on degraded reefs: rather than writing off bleached equatorial reefs as lost causes, the high ROI values in these regions suggest they have a high latent biodiversity potential. If stress-tolerant host genotypes can be successfully outplanted to these Ghost Habitat hotspots, the climate conditions are already setup to support a diverse fish community. This reframes the conservation challenge from passive preservation to active ecosystem engineering, where the restoration of the biotic partner could be key future factor for the climatic potential of the reef Hoegh-Guldberg et al. (2008).

### 4.6 Limitations and Future Directions

While these findings provide a robust mechanism for future range loss, the SDM approach has inherent limitations that frame the interpretation of these results. firstly, similar to many broad-scale ecological studies Breiner et al. (2015); Liu et al. (2024), our occurrence data was sourced from aggregated databases (GBIF, OBIS). Such datasets can contain geographical biases in sampling effort. Given the challenges of direct observation, emerging techniques such as environmental DNA (eDNA) metabarcoding could be valuable for detecting the presence of anemonefish and their specific hosts within potential new habitats or deep-water refugia that visual surveys miss Açıkba et al. (2024). Secondly, the models assume static host-association patterns. While host specificity is generally rigid in anemonefish Fautin and Allen (1997); Litsios et al. (2012), extreme climate stress could theoretically induce host-switching or plasticity that is not captured here. Our models treat species as fixed ecological units, yet local adaptation may allow some populations to persist in conditions deemed unsuitable by the global model. Future work should aim to integrate genomic data to explore intraspecific variation in host use, as cryptic generalism could buffer some populations against the predicted range lags Donelson et al. (2012). Thirdly, while we identify potential range shifts based on climatic and biotic suitability, we did not explicitly model the biophysical connectivity required to realise these shifts. Anemonefish rely on larval dispersal to colonise new reefs. Integrating biophysical dispersal models with these SDMs would likely reveal even more severe constraints, as ocean currents may not align with the necessary poleward trajectories, creating oceanographic barriers that further fragment the Ghost Habitat Treml et al. (2008). Fourthly, existing research indicates that widespread anemonefish and host species often comprise of multiple evolutionary significant units (ESUs) or cryptic species with distinct thermal tolerances Nuryanto and Kochzius (2009). By modelling at the nominal species level, we may have overestimated the niche breadth of certain taxa. If what appears to be a broad-niche generalist is actually a complex of narrow-niche cryptic species, the community vulnerability to climate change may be significantly higher than reported here. finally, an important limitation is that our biotic predictor accounts for host *presence*, but not host *condition*. The SDMs predict where hosts can exist, but not whether they will be healthy. Host sea anemones are susceptible to bleaching and size reduction under thermal stress, which can lead to reproductive failure in the associated fish even if the host survives Hobbs et al. (2013). Consequently, our estimates of suitable habitat are likely optimistic; the realised niche may be even smaller if thermally stressed hosts are physiologically incapable of sustaining symbionts.

Based on these considerations, future research should aim to:

1. Collect targeted, field-verified occurrence data including true absence records to refine model calibration.
2. Integrate population-level genetic data to model distinct ESUs and their specific environmental requirements.
3. Couple SDMs with biophysical models of larval dispersal to generate dynamically realistic projections of recruitment.
4. Investigate the potential for plasticity in host use and the adaptive capacity of both anemonefish and host anemones through experimental evolution studies.
5. Explicitly model changes in interspecific interactions, such as competition for hosts, which may intensify as species’ ranges shift and novel communities assemble Blois et al. (2013).

Addressing these limitations will transition the field from predicting potential distributions to forecasting population persistence, providing the robust, mechanistic tools necessary for effective coral reef conservation.

## 5. CONCLUSION

This study demonstrates that for obligate mutualists, climate change is not just a physiological challenge but a relational one. By removing the assumption that biotic interactions are negligible at macro-ecological scales, we have shown that the future distribution of anemonefish will be defined not by where they can survive, but by where their partners can persist. The finding of widespread Ghost Habitats shows that without accounting for the immobile half of the mutualism, we are systematically underestimating the extinction risk of the mobile half. Our findings show that species with narrow environmental tolerances are trapped facing the highest climate velocities while also being tethered to the most dispersal-limited hosts. In contrast, generalism offers only a conditional buffer, effective only as long as the most resilient host in the assemblage persists. Ultimately, these results argue for a shift in coral reef conservation. We must move beyond single-species management and towards a framework that prioritises the resilience of the interaction itself.

## Abbreviations

CPI: Conservation Priority Index
MEOW: Marine Ecoregions of the World
MSP: Marine Spatial Planning
ROI: Restoration Opportunity Index
SDM: Species Distribution Model
SSP: Shared Socioeconomic Pathway

## acknowledgements

I would like express his thanks to all the teaching staffs in the Kyoto University Department of Information Science for their contribution.

