## Supplementary Material for "Effects of Host-Dependent Niches and Biotic Constraints on Climate Change Driven Range Shifts in Anemonefish"

#### Supporting Information

#### Data and Code Availability

The complete reproducible analysis workflow is openly available.

##### | Source Code & Data

Full repository with all scripts for data processing and modelling is available on GitHub at: `https://github.com/chrisrauch193/sdm_anemonefish`

##### | Interactive Results & figures

High-resolution supplementary figures, maps, and rendered post-analysis report viewable at: `https://chrisrauch193.github.io/sdm_anemonefish/`

#### S1. Environmental Variable Selection

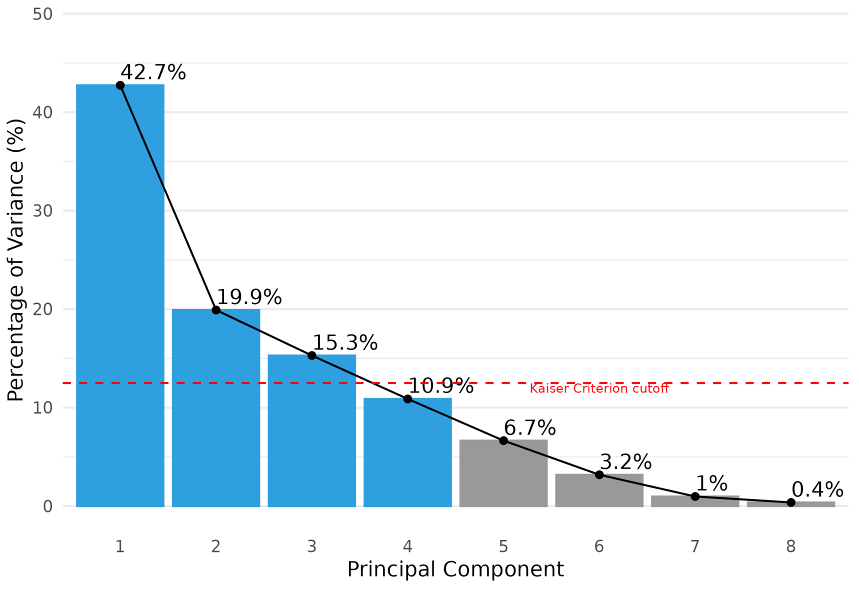


**FIGURE S1** Scree plot from the Principal Component Analysis (PCA) of eight bioclimatic variables. The blue bars indicate the four components (PC1–PC4) retained for modeling, which together explain 88.8% of the total variance. The red dashed line indicates the Kaiser-Guttman criterion (Eigenvalue = 1).

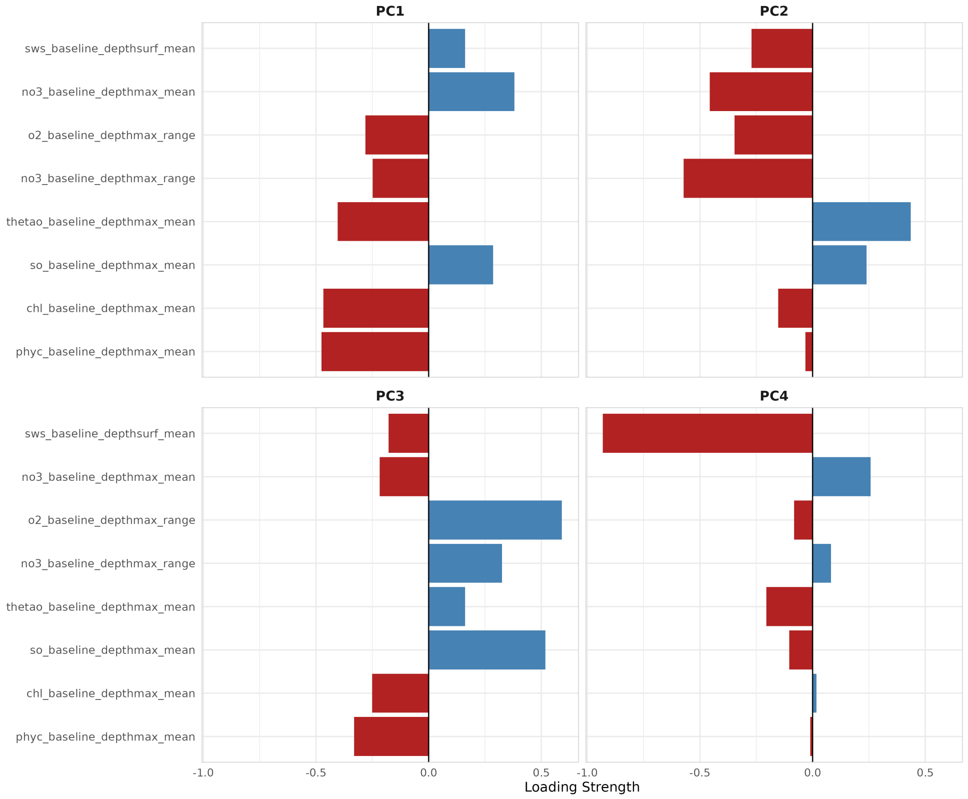


**FIGURE S2** Factor loadings for the four retained Principal Components. Bars indicate the strength and direction of contribution for each original bioclimatic variable to the new synthetic axes. PC1 represents a primary gradient of surface temperature and light (PAR), while PC2 captures gradients in nutrients (Nitrate) and depth-specific temperature ranges.

#### S2. Supplementary Tables

**TABLE S1** Extended model performance metrics for the Combined Species Distribution Models (SDMs) of all 17 analysed anemonefish species. Values represent the mean ± standard deviation across bootstrap iterations. CBI = Continuous Boyce Index.

| Species | AUC | TSS | CBI | Brier Score | LogLoss |
|---|---|---|---|---|---|
| *Amphiprion akallopisos* | 0.721 ± 0.109 | 0.446 ± 0.176 | 0.095 ± 0.354 | 0.133 ± 0.039 | 0.402 ± 0.092 |
| *Amphiprion akindynos* | 0.927 ± 0.042 | 0.779 ± 0.114 | 0.819 ± 0.165 | 0.067 ± 0.032 | 0.231 ± 0.088 |
| *Amphiprion allardi* | 0.886 ± 0.101 | 0.759 ± 0.174 | 0.528 ± 0.419 | 0.075 ± 0.052 | 0.252 ± 0.162 |
| *Amphiprion chrysogaster* | 0.815 ± 0.024 | 0.676 ± 0.030 | 0.730 ± 0.137 | 0.077 ± 0.007 | 0.276 ± 0.020 |
| *Amphiprion chrysopterus* | 0.927 ± 0.014 | 0.748 ± 0.032 | 0.852 ± 0.081 | 0.051 ± 0.008 | 0.179 ± 0.025 |
| *Amphiprion ephippium* | 0.908 ± 0.033 | 0.770 ± 0.085 | 0.718 ± 0.198 | 0.083 ± 0.022 | 0.281 ± 0.063 |
| *Amphiprion frenatus* | 0.815 ± 0.077 | 0.554 ± 0.139 | 0.491 ± 0.283 | 0.113 ± 0.026 | 0.358 ± 0.072 |
| *Amphiprion fuscocaudatus* | 0.803 ± 0.061 | 0.607 ± 0.138 | 0.537 ± 0.252 | 0.093 ± 0.022 | 0.308 ± 0.063 |
| *Amphiprion latifasciatus* | 0.765 ± 0.201 | 0.642 ± 0.242 | 0.279 ± 0.641 | 0.073 ± 0.046 | 0.245 ± 0.135 |
| *Amphiprion leucokranos* | 0.942 ± 0.029 | 0.861 ± 0.079 | 0.731 ± 0.237 | 0.038 ± 0.017 | 0.137 ± 0.052 |
| *Amphiprion melanopus* | 0.829 ± 0.080 | 0.582 ± 0.157 | 0.522 ± 0.273 | 0.108 ± 0.028 | 0.345 ± 0.080 |
| *Amphiprion nigripes* | 0.794 ± 0.202 | 0.558 ± 0.242 | 0.366 ± 0.531 | 0.085 ± 0.048 | 0.279 ± 0.128 |
| *Amphiprion ocellaris* | 0.835 ± 0.090 | 0.623 ± 0.151 | 0.621 ± 0.219 | 0.107 ± 0.025 | 0.339 ± 0.070 |
| *Amphiprion percula* | 0.914 ± 0.015 | 0.715 ± 0.029 | 0.865 ± 0.054 | 0.063 ± 0.007 | 0.220 ± 0.021 |
| *Amphiprion polymnus* | 0.911 ± 0.020 | 0.733 ± 0.050 | 0.817 ± 0.091 | 0.066 ± 0.010 | 0.225 ± 0.028 |
| *Amphiprion rubrocinctus* | 0.977 ± 0.011 | 0.918 ± 0.023 | 0.925 ± 0.073 | 0.030 ± 0.008 | 0.114 ± 0.025 |
| *Amphiprion tricinctus* | 0.965 ± 0.023 | 0.879 ± 0.051 | 0.794 ± 0.245 | 0.039 ± 0.015 | 0.138 ± 0.046 |

**TABLE S2** Model performance metrics for the eight Host Sea Anemone species (EnvOnly models).

| Species | AUC | TSS | CBI | Brier |
|---|---|---|---|---|
| *Cryptodendrum adhaesivum* | 0.809 ± 0.012 | 0.525 ± 0.023 | 0.556 ± 0.210 | 0.104 ± 0.015 |
| *Entacmaea quadricolor* | 0.755 ± 0.021 | 0.416 ± 0.035 | 0.864 ± 0.050 | 0.096 ± 0.012 |
| *Heteractis aurora* | 0.745 ± 0.015 | 0.401 ± 0.028 | 0.459 ± 0.180 | 0.094 ± 0.010 |
| *Heteractis crispa* | 0.790 ± 0.018 | 0.541 ± 0.041 | 0.720 ± 0.150 | 0.089 ± 0.018 |
| *Heteractis magnifica* | 0.767 ± 0.020 | 0.462 ± 0.033 | 0.580 ± 0.190 | 0.091 ± 0.012 |
| *Heteractis malu* | 0.714 ± 0.033 | 0.455 ± 0.061 | 0.350 ± 0.250 | 0.105 ± 0.020 |
| *Stichodactyla gigantea* | 0.730 ± 0.025 | 0.425 ± 0.040 | 0.410 ± 0.220 | 0.098 ± 0.015 |
| *Stichodactyla haddoni* | 0.735 ± 0.019 | 0.407 ± 0.031 | 0.480 ± 0.180 | 0.095 ± 0.012 |

**TABLE S3** Projected poleward range shift (km) for all anemonefish species under the high-emission scenario (SSP5-8.5, 2100). Shifts are calculated as the distance between the current and future centroids of the Combined model suitability.

| Species | Group | Range Shift (km) |
|---|---|---|
| *Amphiprion nigripes* | Specialist | 2881.3 |
| *Amphiprion akindynos* | Generalist | 2658.3 |
| *Amphiprion chrysogaster* | Generalist | 2172.5 |
| *Amphiprion latifasciatus* | Generalist | 2017.2 |
| *Amphiprion fuscocaudatus* | Generalist | 1633.8 |
| *Amphiprion rubrocinctus* | Specialist | 1579.0 |
| *Amphiprion akallopisos* | Specialist | 1485.6 |
| *Amphiprion allardi* | Generalist | 1360.1 |
| *Amphiprion ephippium* | Specialist | 1353.8 |
| *Amphiprion percula* | Specialist | 965.5 |
| *Amphiprion melanopus* | Specialist | 838.4 |
| *Amphiprion polymnus* | Generalist | 617.7 |
| *Amphiprion frenatus* | Specialist | 547.2 |
| *Amphiprion leucokranos* | Generalist | 519.9 |
| *Amphiprion tricinctus* | Generalist | 337.0 |
| *Amphiprion ocellaris* | Specialist | 70.2 |

**TABLE S4** final number of spatially unique occurrence records (*N*) used for model training after spatial thinning and quality control.

| Species | final Occurrence Count (*N*) |
|---|---|
| **Anemonefish** | |
| *Amphiprion akallopisos* | 230 |
| *Amphiprion akindynos* | 374 |
| *Amphiprion allardi* | 123 |
| *Amphiprion chrysogaster* | 46 |
| *Amphiprion chrysopterus* | 415 |
| *Amphiprion ephippium* | 41 |
| *Amphiprion frenatus* | 320 |
| *Amphiprion fuscocaudatus* | 23 |
| *Amphiprion latifasciatus* | 47 |
| *Amphiprion leucokranos* | 26 |
| *Amphiprion melanopus* | 634 |
| *Amphiprion nigripes* | 163 |
| *Amphiprion ocellaris* | 790 |
| *Amphiprion percula* | 275 |
| *Amphiprion polymnus* | 182 |
| *Amphiprion rubrocinctus* | 93 |
| *Amphiprion tricinctus* | 34 |
| **Host Sea Anemones** | |
| *Cryptodendrum adhaesivum* | 192 |
| *Entacmaea quadricolor* | 856 |
| *Heteractis aurora* | 270 |
| *Heteractis crispa* | 136 |
| *Heteractis magnifica* | 249 |
| *Heteractis malu* | 30 |
| *Stichodactyla gigantea* | 178 |
| *Stichodactyla haddoni* | 212 |

### S3. Supplementary Maps

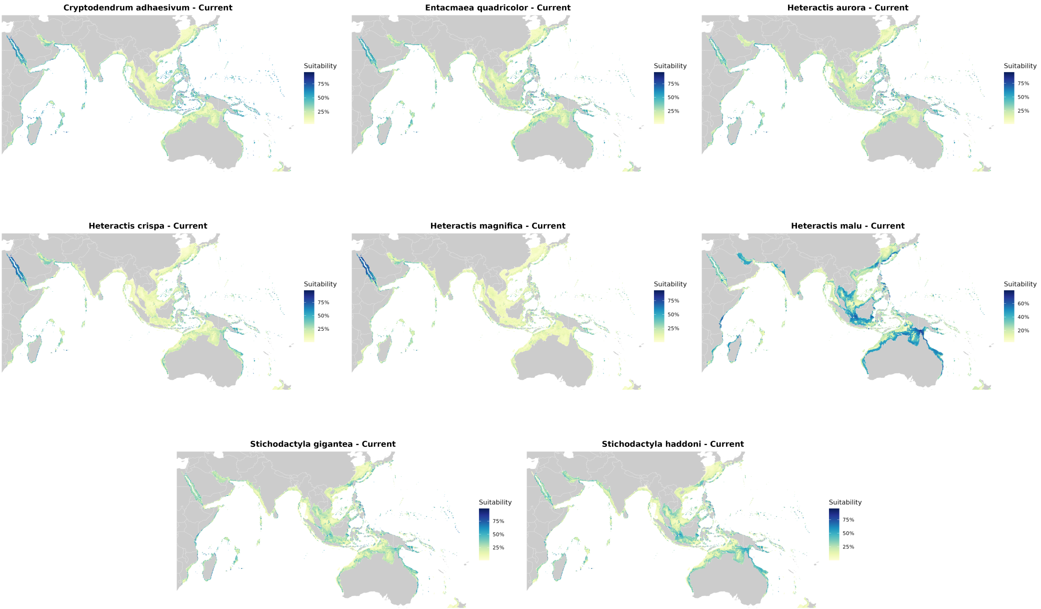


**FIGURE S3** Current suitability maps for Host Sea Anemone species (EnvOnly Model).

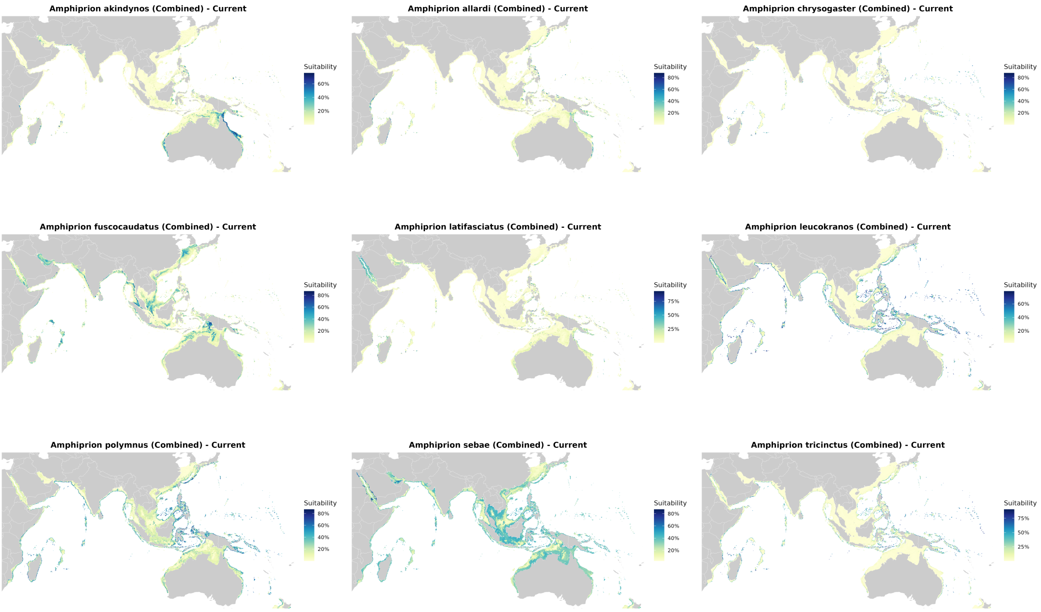


**FIGURE S4** Current suitability maps for generalist anemonefish species (Combined Model).

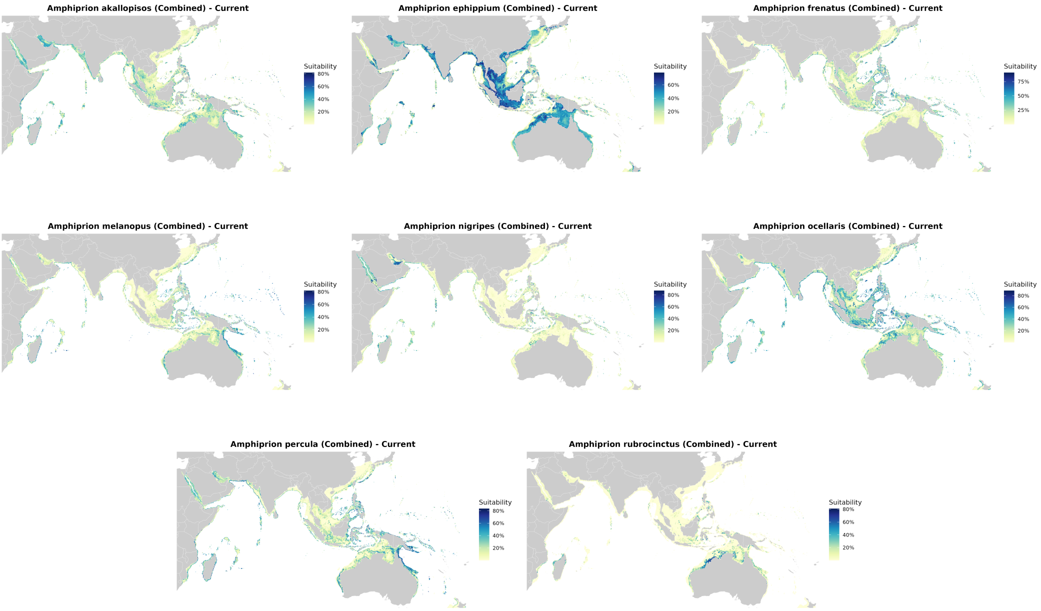


**FIGURE S5** Current suitability maps for specialist anemonefish species (Combined Model).

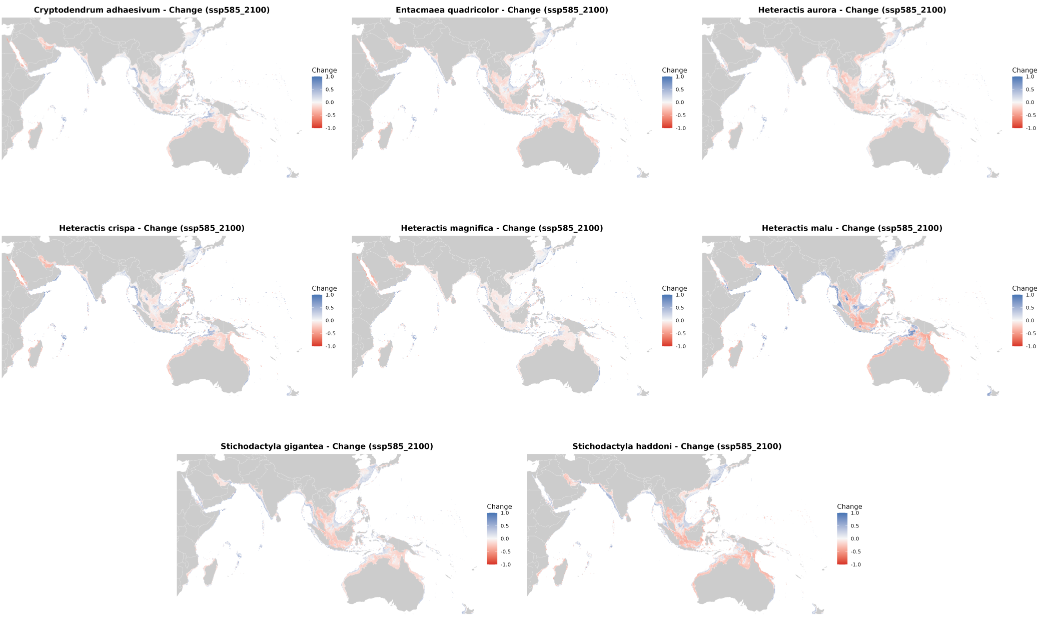


**FIGURE S6** Projected change in suitability (Delta) for Host Sea Anemones under SSP5-8.5 (2100).

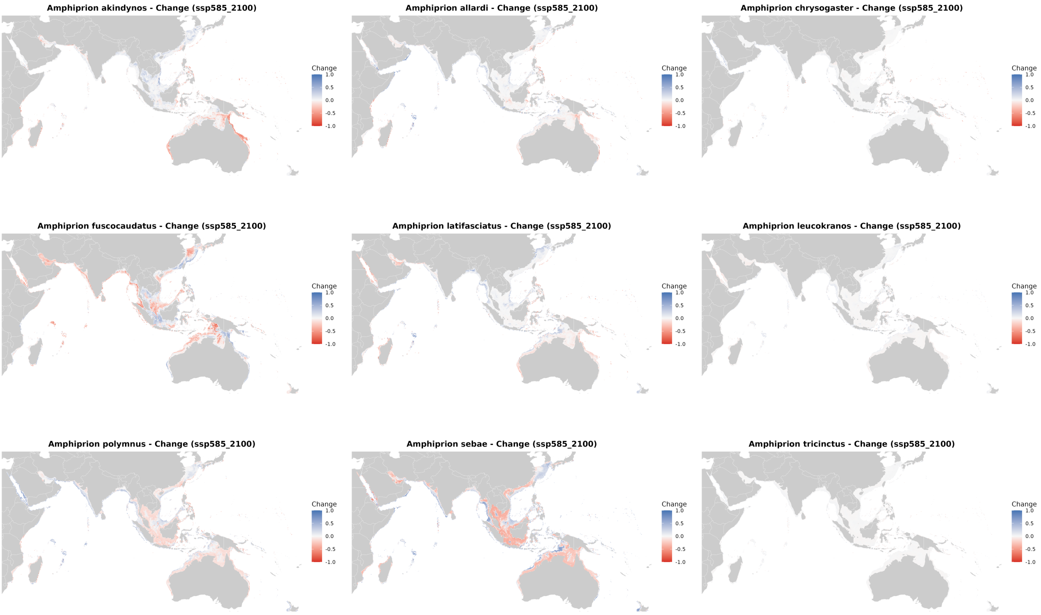


**FIGURE S7** Projected change in suitability (Delta) for Generalist fish under SSP5-8.5 (2100).

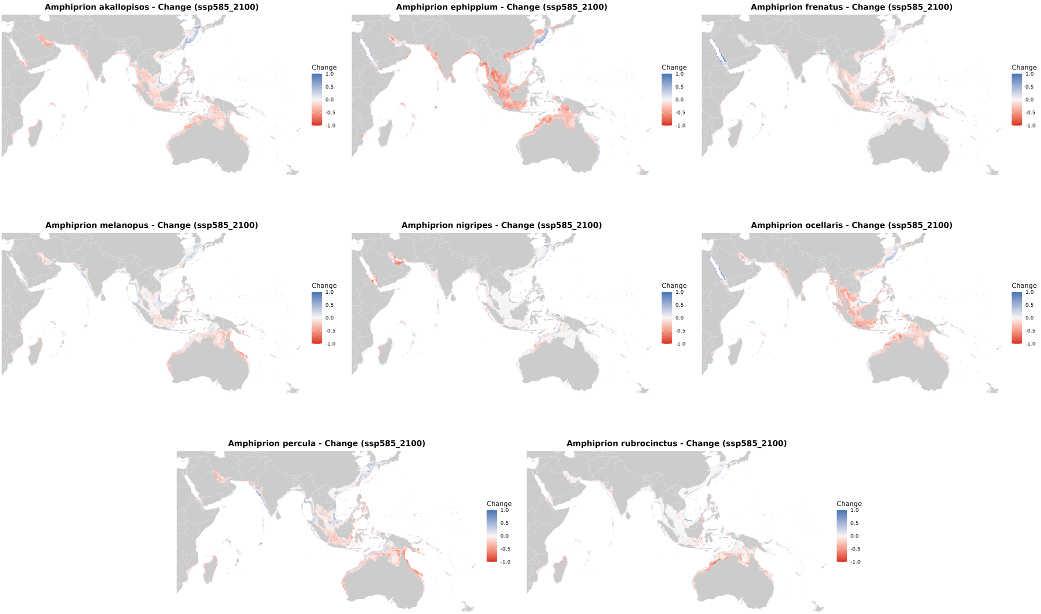


**FIGURE S8** Projected change in suitability (Delta) for Specialist fish under SSP5-8.5 (2100).

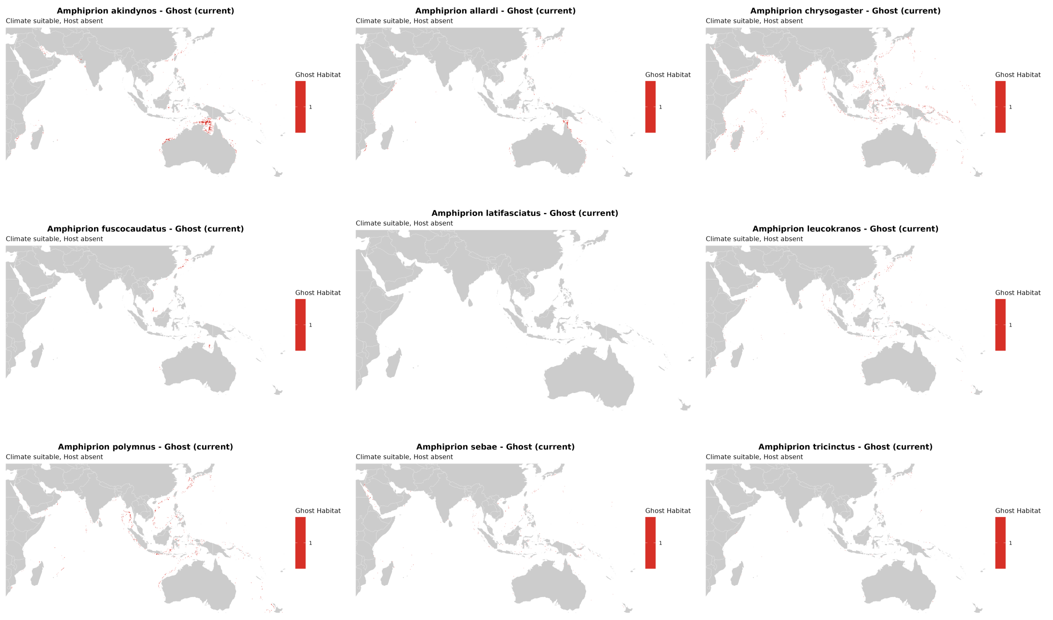


**FIGURE S9** Current Ghost Habitat for generalist fish. Red areas indicate climatically suitable but host-free regions.

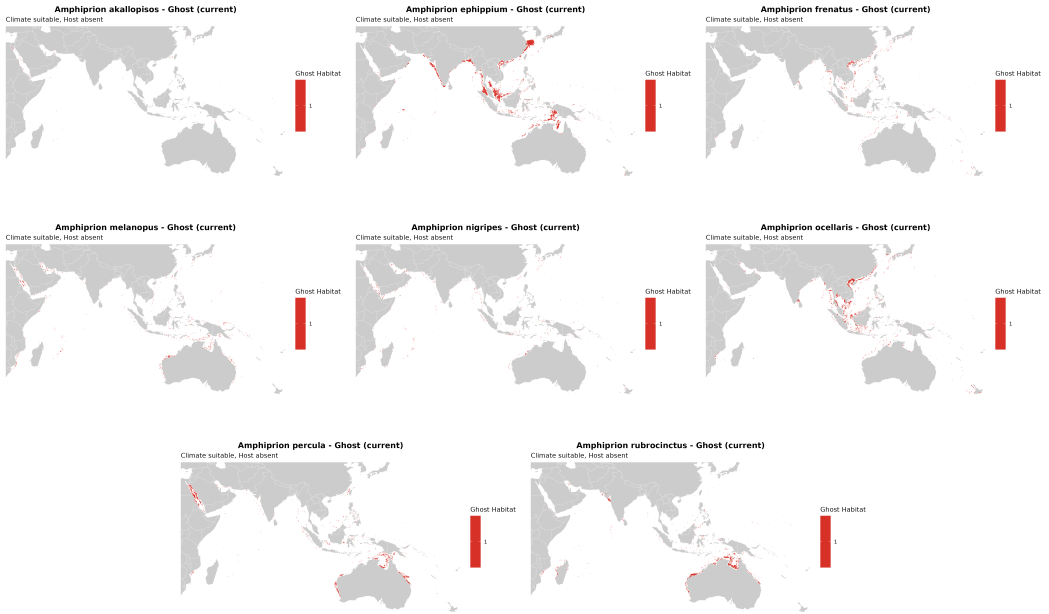


**FIGURE S10** Current Ghost Habitat for specialist fish. Red areas indicate climatically suitable but host-free regions.

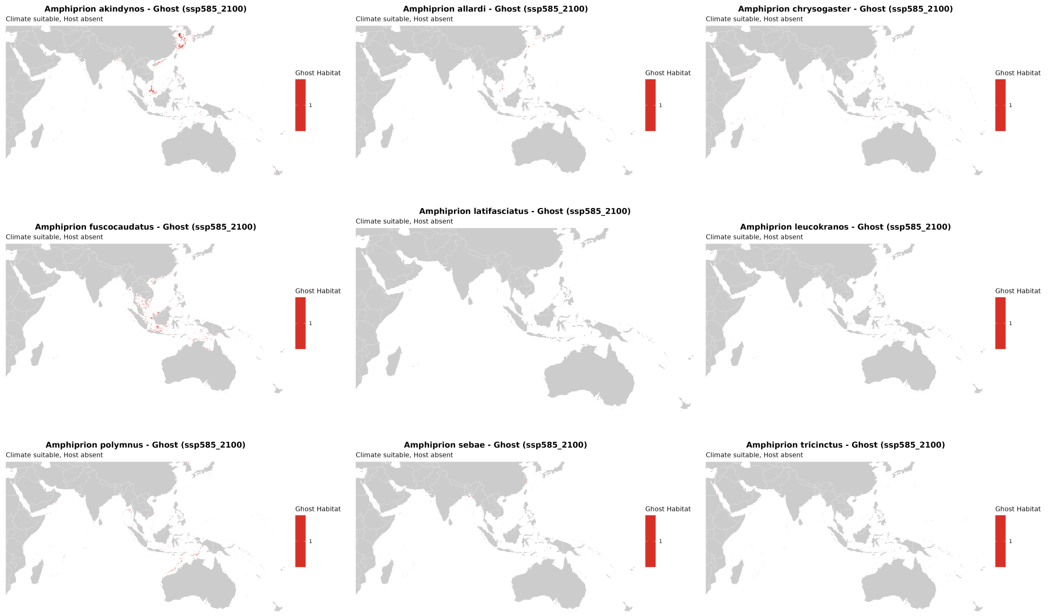


**FIGURE S11** Projected Ghost Habitat for generalist fish under SSP5-8.5 (2100).

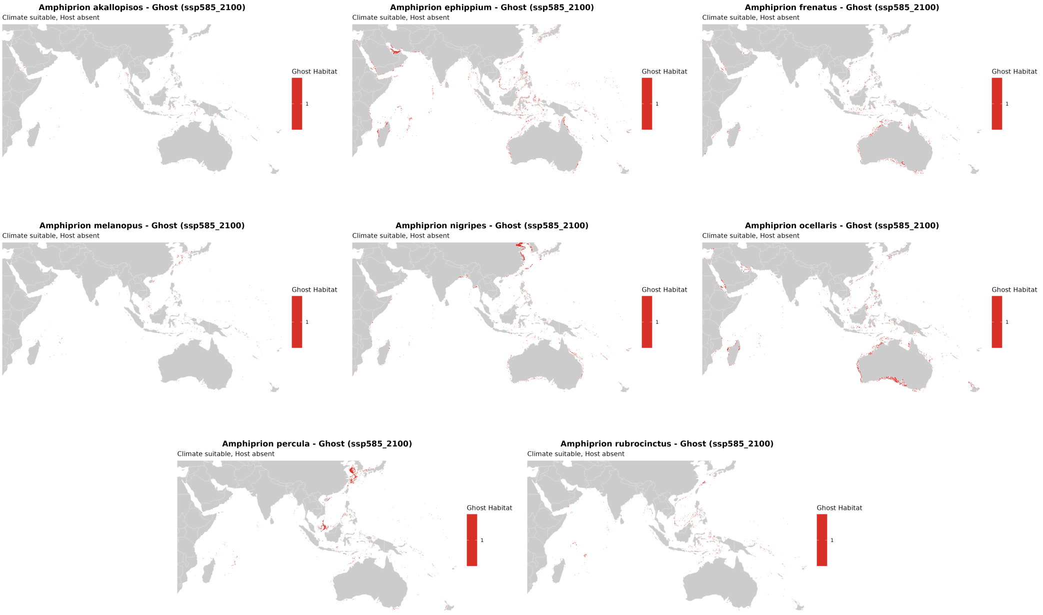


**FIGURE S12** Projected Ghost Habitat for specialist fish under SSP5-8.5 (2100).

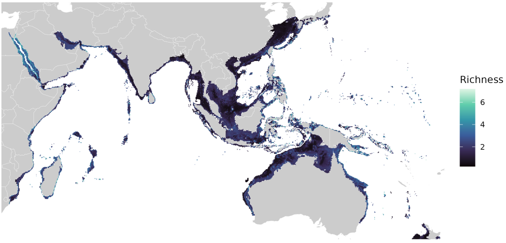


(A) Current Host Richness

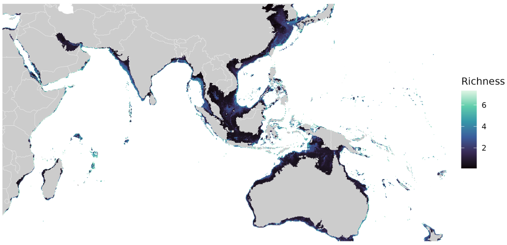


(B) Host Richness (SSP1-1.9 2050)

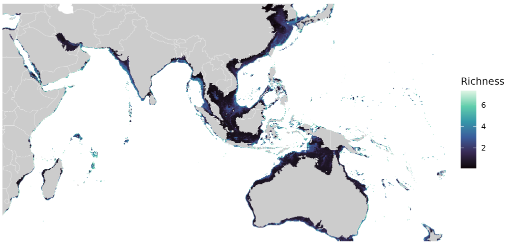


(C) Host Richness (SSP1-1.9 2100)

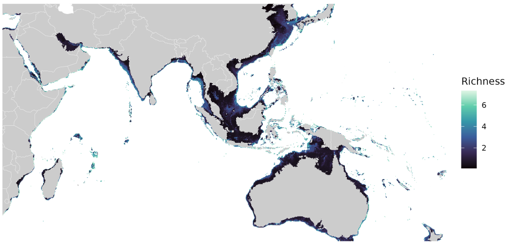


(D) Host Richness (SSP5-8.5 2050)

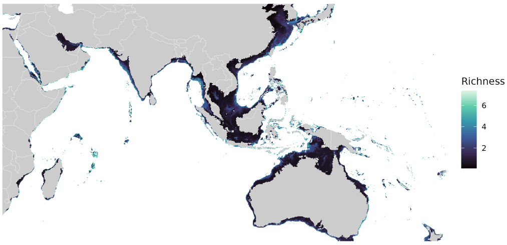


(E) Host Richness (SSP5-8.5 2100)

**FIGURE S13** Projected species richness for the host sea anemone guild across all scenarios. Warmer colors indicate higher biodiversity.

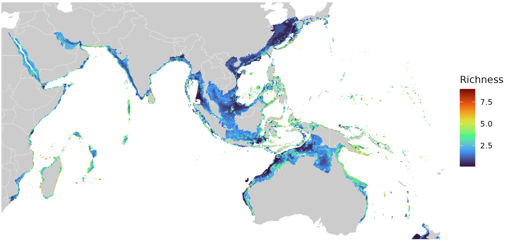


**(A) Current fish Richness**

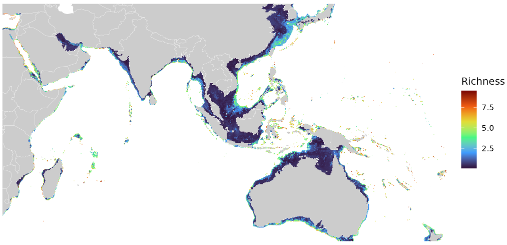


**(B) fish Richness (SSP1-1.9 2050)**

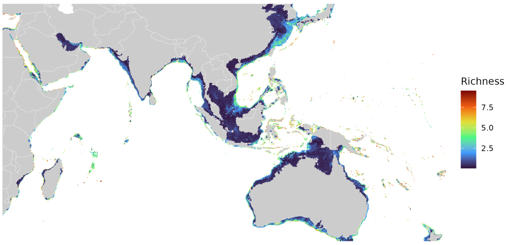


**(C) fish Richness (SSP1-1.9 2100)**

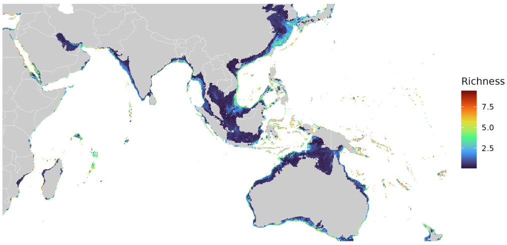


**(D) fish Richness (SSP5-8.5 2050)**

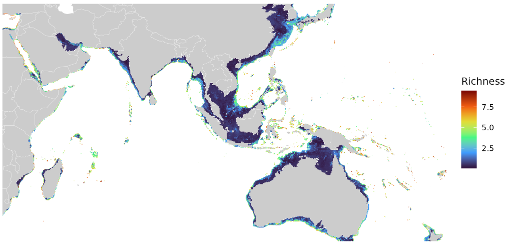


**(E) fish Richness (SSP5-8.5 2100)**

**FIGURE S14** Projected species richness for the anemonefish guild (Combined Models) across all scenarios. Note the contraction of high-richness zones in the equatorial tropics under SSP5-8.5.
